# Comparative molecular landscapes of immature neurons in the mammalian dentate gyrus across species reveal special features in humans

**DOI:** 10.1101/2025.02.16.638557

**Authors:** Yi Zhou, Yijing Su, Qian Yang, Jiaqi Li, Yan Hong, Taosha Gao, Yanqing Zhong, Xueting Ma, Mengmeng Jin, Xinglan Liu, Nini Yuan, Benjamin C. Kennedy, Lizhou Wang, Longying Yan, Angela N. Viaene, Ingo Helbig, Sudha K. Kessler, Joel E. Kleinman, Thomas M. Hyde, David W. Nauen, Cirong Liu, Zhen Liu, Zhiming Shen, Chao Li, Shengjin Xu, Jie He, Daniel R. Weinberger, Guo-li Ming, Hongjun Song

**Affiliations:** Department of Neuroscience and Mahoney Institute for Neurosciences, Perelman School of Medicine, University of Pennsylvania, Philadelphia, PA, USA; Institute of Neuroscience, State Key Laboratory of Neuroscience, CAS Center for Excellence in Brain Science and Intelligence Technology, Chinese Academy of Sciences, Shanghai, China; University of Chinese Academy of Sciences, Shanghai, China; Department of Oral Medicine, School of Dental Medicine, University of Pennsylvania, Philadelphia, PA, USA; Division of Neurosurgery, Children’s Hospital of Philadelphia, Philadelphia, PA, USA.; Department of Neurosurgery, Perelman School of Medicine, University of Pennsylvania, Philadelphia, PA, USA; Department of Pathology and Laboratory Medicine, Children’s Hospital of Philadelphia, Philadelphia, PA, USA; Division of Neurology, Children’s Hospital of Philadelphia, Philadelphia, PA, USA; The Epilepsy NeuroGenetics Initiative (ENGIN), Children’s Hospital of Philadelphia, Philadelphia, PA, USA; Department of Biomedical and Health Informatics (DBHi), Children’s Hospital of Philadelphia, Philadelphia, PA, USA; Department of Neurology, Perelman School of Medicine, University of Pennsylvania, Philadelphia, PA, USA; Lieber Institute for Brain Development, The Solomon H. Snyder Department of Neuroscience, Department of Neurology, and Department of Psychiatry, School of Medicine, Johns Hopkins University, Baltimore, MD, USA; Department of Pathology, Johns Hopkins University School of Medicine, Baltimore, MD, USA.; Department of Cell and Developmental Biology, Perelman School of Medicine, University of Pennsylvania, Philadelphia, PA, USA; Institute for Regenerative Medicine, University of Pennsylvania, Philadelphia, PA, USA.; Department of Psychiatry, Perelman School of Medicine, University of Pennsylvania, Philadelphia, PA, USA; The Epigenetics Institute, Perelman School of Medicine, University of Pennsylvania, Philadelphia, PA, USA

**Keywords:** immature neuron, hippocampus, dentate gyrus, adult neurogenesis, evolution, cross-species comparison, human-specific feature, non-human primate, single-cell RNA sequencing, machine learning

## Abstract

Immature dentate granule cells (imGCs) arising from adult hippocampal neurogenesis contribute to plasticity, learning and memory, but their evolutionary changes across species and specialized features in humans remain poorly understood. Here we performed machine learning-augmented analysis of published single-cell RNA-sequencing datasets and identified macaque imGCs with transcriptome-wide immature neuronal characteristics. Our cross-species comparisons among humans, monkeys, pigs, and mice showed few shared (such as DPYSL5), but mostly species-specific gene expression in imGCs that converged onto common biological processes regulating neuronal development. We further identified human-specific transcriptomic features of imGCs and demonstrated functional roles of human imGC-enriched expression of a family of proton-transporting vacuolar-type ATPase subtypes in development of imGCs derived from human pluripotent stem cells. Our study reveals divergent gene expression patterns but convergent biological processes in the molecular characteristics of imGCs across species, highlighting the importance of conducting independent molecular and functional analyses for adult neurogenesis in different species.

## Main Text

Adult hippocampal neurogenesis occurs throughout life in most mammals examined and its dysregulation has been implicated in various neurological disorders^1–7^. The functional outcomes of adult hippocampal neurogenesis are primarily mediated by immature dentate granule cells (imGCs), which exhibit different molecular, cellular, and physiological properties compared to mature dentate granule cells (mGCs)^8–11^. There have been extensive investigations of imGCs in the hippocampus using mouse models^4,12–15^, including transcriptomic profiling using single-cell or single-nucleus RNA sequencing (scRNA-seq)^16–24^. Recent studies have extended the application of scRNA-seq analyses to explore molecular characteristics of imGCs in other species, especially in non-human primates^25–30^ and humans^27–35^, where there appears to be substantial gene expression differences between imGCs of humans and mice, yet the extent to which these features are specialized in humans remains unclear^30,31,36–38^. Interspecies divergence in mammalian brains has been linked to remarkable capabilities of humans compared to other species^39–41^. Emerging cross-species comparative analyses of the developing cortex have identified variations in individual genes or regulatory elements that contribute to distinct cellular and physiological features in humans and non-human primates, particularly during the neuronal maturation process^42–44^. Identifying molecular features uniquely enriched in human hippocampal imGCs holds promise for elucidating their specialized molecular regulation during human neuronal development and function. Moreover, as most knowledge stems from mouse models, it is imperative to delve deeper into the understanding of species-specific differences to select the most suitable research models when developing therapeutic strategies to regulate adult hippocampal neurogenesis in patients^37^. However, a systematic comparison of the molecular landscapes of immature neurons in the hippocampus across species is lacking due to inconsistent cell type identification in humans and macaques, impeding further molecular analysis^1,30,37,45^.

Most studies that identify imGCs in non-mouse species rely on the molecular signatures of mouse imGCs - they either identify imGCs using conventional markers obtained from mouse studies or they apply transfer learning from mouse imGC clusters to annotate cell clusters in scRNA-seq datasets of other species^30^. So far, both marker-based histological analyses^27,46–48^ and scRNA-seq analyses^25–28,35^ showed inconsistent findings regarding the identities and abundance of imGCs in the postnatal hippocampus of humans and macaques^30,49,50^. In particular, previous scRNA-seq studies of the postnatal macaque hippocampus that aimed to identify immature neuronal clusters based on unsupervised learning showed results either supporting^25–27^ or refuting^28^ the presence of immature neurons (**Supplementary Table 1**). For example, several studies^25–27^ used unsupervised clustering followed by annotation of imGCs based on one or two predefined markers that were identified in mice. One study^25^ marked a cluster as imGCs that was enriched for PROX1 and SNAP25, which are common gene markers for all dentate granule cells (GCs) instead of imGCs. Another study^26^ marked distinct imGC clusters that contained 7,547 cells out of 34,039 GCs (22.2%) in young adult (4-6 years) and middle-aged adult macaques (13-15 years), a proportion significantly higher than the histological detection of imGCs in age-matched macaques^27,46,47^. They also did not observe any differences in the percentages of imGCs among all GCs between the two age groups^26^. We performed our own analysis of this dataset^26^. By scoring^51^ their similarities to a gene module^16^ of mouse immature neuronal progeny and examining their associated Gene Ontology (GO) terms of biological processes, we found that the genes enriched in the annotated imGC clusters in this study^26^ displayed very limited immature neuronal characteristics (**Extended Data Fig. 1a, b**). A third study^28^ used transfer learning^52,53^ to identify neuronal progenitors in adult macaques using mouse datasets, but they did not examine the existence or abundance of imGCs. Overall, these findings raise a question about the general assumption that imGCs must share abundant molecular similarities across different mammalian species^30^ and highlight the need for systematic cross-species comparative analyses of imGCs across studies using a consistent approach to examine their conserved and divergent features at the whole transcriptome level.

Here, we provide a holistic view of the comparative molecular landscapes of imGCs in the hippocampus of four different mammalian species, including humans, macaques, pigs, and mice, using published datasets^16,25–29,31,34^ (**Supplementary Table 1**). We first identified the imGC population that exhibits immature neuronal characteristics in published scRNA-seq datasets of the macaque hippocampus using a machine learning-augmented approach, which was previously validated for efficacy and specificity in identifying human and mouse imGCs^31^. Next, we performed cross-species comparisons of imGCs among published scRNA-seq datasets of humans, macaques, pigs, and mice and we discovered substantial molecular differences across species and many human-enriched features, shedding light on their specialized functions and regulation unique to human imGCs. Finally, we examined the functional roles of one human imGC-enriched gene family in regulating development of imGCs-derived from human induced pluripotent stem cells (iPSCs).

## Results

### Identification of macaque imGCs across datasets by machine learning

Prior to meta-analyses of hippocampal scRNA-seq datasets of multiple mammalian species, we first aimed to identify imGCs with immature neuronal characteristics across all available macaque hippocampal scRNA-seq datasets from different studies^25–28^. Given the potential variations in the sample preparation and qualities of annotation and alignment of transcriptomes across different species, we refrained from utilizing classifiers developed for mice in the context of macaques. Instead, we employed a validated, prototype-based, supervised learning method trained with macaque datasets to distinguish macaque imGCs from mGCs (**Fig. 1a**), following a similar strategy that we had previously established for human and mouse datasets^31^. Specifically, we used high-confidence DCX^+^PROX1^+^CALB1^-^ imGCs selected from the GC cluster in a published young macaque hippocampal dataset (4-6 years)^25^ as cell prototypes to train a machine learning classifier, where macaque imGCs are defined by a transcriptome-wide weighted panel of genes (**Fig. 1a; Extended Data Fig. 2a, b; Supplementary Table 2;** see **Methods**). The positively weighted genes defining the macaque imGC classifier, similar to those of humans and mice^31^, showed GO signatures related to neurogenesis, neurite development, and synaptic plasticity, indicating a convergence in neuron development and plasticity gene programs in imGCs of all three species (**Fig. 1b, c; Supplementary Table 3**). Surprisingly, individual weighted genes involved in these programs are highly divergent, even between humans and macaques (**Fig. 1d; Extended Data Fig. 2c**).

**Figure 1.**
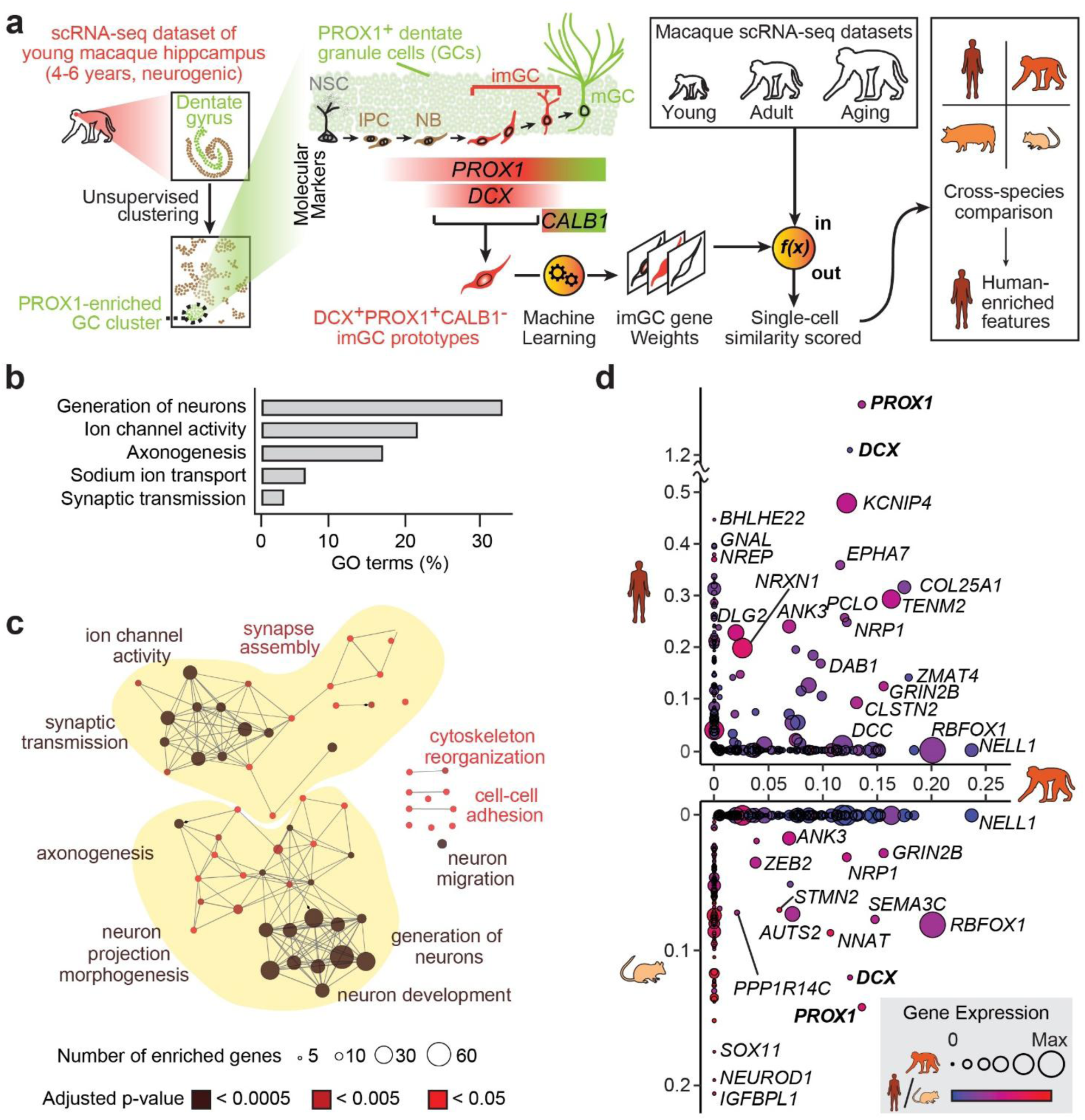
Machine learning-augmented identification and molecular characteristics of immature neurons in macaque hippocampal single-cell RNA sequencing datasets. **a,** Schematic illustration of the experimental design. scRNA-seq, single-cell or single-nucleus RNA sequencing; NSC, neural stem cell; IPC, intermediate progenitor cell; NB, neuroblast; GC, dentate granule cell; imGC, immature GC; mGC, mature GC. **b, c,** Top Gene Ontology (GO) term groups (**b**) and GO networks (**c**) of biological processes of the positive gene weights defining macaque imGCs (using annotations based on our machine learning model), coloured by false discovery rate (FDR)-adjusted p-value (c). Only significantly enriched nodes are displayed (one-sided hypergeometric test, FDR-adjusted *P* < 0.05) (c). The node size represents the term enrichment significance. Examples of the most significant terms per group are shown (**c**). **d,** Comparison of positive gene weights defining imGCs in humans, macaques, and mice generated by independent machine learning models for each species, shown on the x-axis (left), y-axis, and x-axis (right), respectively. The machine learning model for macaque imGCs was generated in this study; models for human and mouse imGCs were previously generated^31^. Dot size represents gene expression in macaque imGCs; dot colour shows gene expression in humans or mouse imGCs in the corresponding plots.

Next, we applied the trained classifier to score each cell in all currently publicly available macaque hippocampal scRNA-seq datasets of various ages and sequencing platforms^25–28^ based on its similarity to prototype imGCs (**Fig. 1a**). All immature neurons identified by this classifier reside within the GC cluster, while none were selected from cells in the widely acknowledged “non-neurogenic” brain regions^2^, including the postnatal *cornu ammonis*, subiculum, entorhinal cortex, or a scRNA-seq dataset of the adult macaque cortex^54^, indicating the specificity of our method (**Fig. 2a; Extended Data Fig. 3a, b**). We observed a decrease in the abundance of imGCs among GCs with age (**Fig. 2b**); the percentage of imGCs among GCs that we identified across ages *in silico* (mean = 5.2 ±2.3 %; median = 1.1 %) and the level of decrease over age are comparable to those analyses using immunohistology and/or nucleotide analogue tracing measurement of newborn neuron levels^47,55,56^ despite a considerable level of variability among samples, providing further validation of our approach. Together, our prototype-based machine learning classifier effectively and specifically identified imGCs in macaque scRNA-seq datasets.

**Figure 2.**
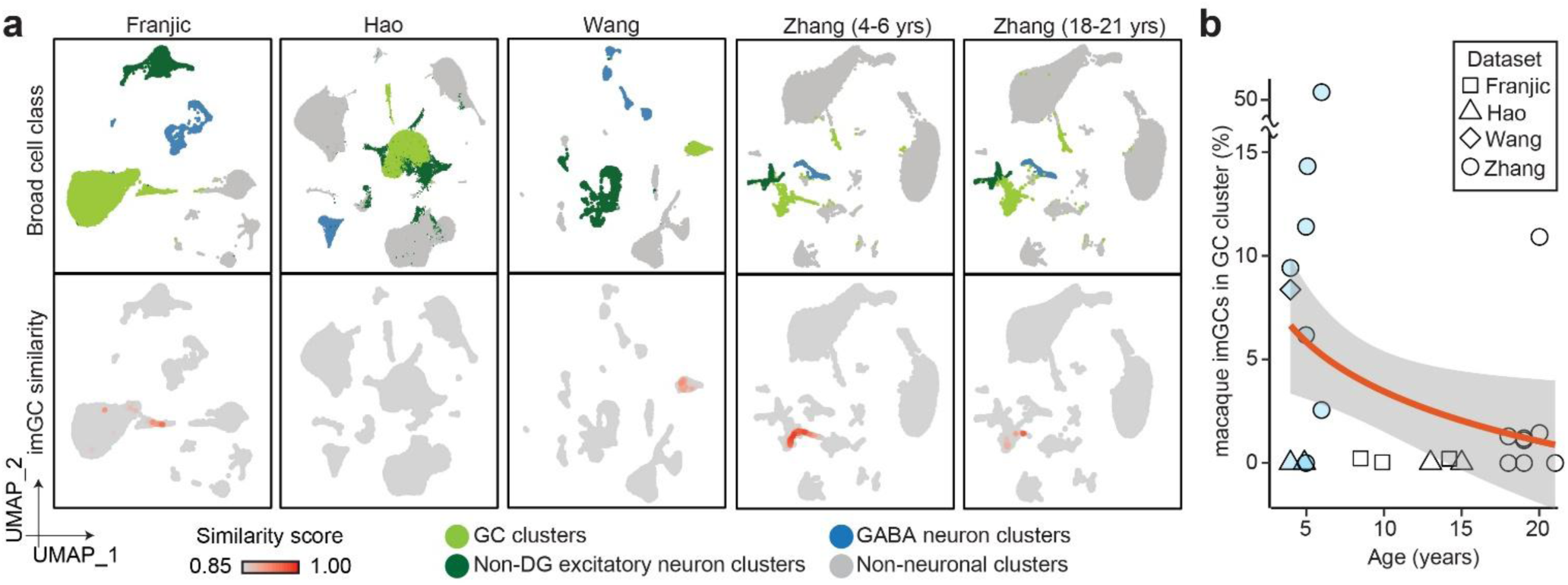
Identification of immature neurons in the postnatal macaque hippocampus across ages and studies. **a,** UMAP plots showing all publicly available scRNA-seq datasets of the macaque hippocampus^25–28^ coloured by four broad cell classes (top rows) and by similarity score to prototypical imGCs (bottom rows). Datasets containing multiple specimens from each study (noted by first author’s last name) were integrated and are shown in aggregate. Note that imGCs are only present in the GC cluster. **b,** Quantification of percentages of imGCs (with similarity score *p* ≥ 0.85) among all GCs in each macaque hippocampal specimen across ages. Data point shapes match published macaque hippocampal datasets^25–28^. Cyan points indicate young macaque group (4-6 years); hollow points indicate adult and aging groups (8-15 and 18-21 years, respectively). Data points are fitted with generalized linear model fitting (orange line) and 95% confidence intervals (light grey shaded areas).

### Divergent imGC-enriched gene expression with convergence in biological processes across species

We next performed cross-species comparative analysis by investigating the conserved and divergent molecular features enriched in imGCs compared to mGCs in published hippocampal scRNA-seq datasets of humans, macaques, pigs, and mice^16,25,28,31,34^ (**Fig. 1a**; see **Methods**). Due to the potential limitations in sequencing coverage inherent to scRNA-seq technology, our analysis may have predominantly captured molecular features of highly expressed genes while neglecting alternative splicing. For the purpose of transcriptome comparisons, we restricted our analysis to genes with orthologs to humans. Given that previous marker-based calling or transfer learning methods using molecular signatures of mouse imGCs had variable success in identifying imGC populations in humans and macaques^26,28,30^, we first evaluated^51^ whether imGCs of different species, including the macaque imGCs identified by our machine learning approach, have immature neuronal characteristics by performing gene module analysis with genes enriched in mouse immature progeny (neuroblasts and imGCs) and mGCs^16^. Human imGCs were identified in our previous study^31^ and macaque imGCs were identified in the current study using their corresponding machine learning models; for pig^28^ and mouse^16^ imGCs, we used annotations based on unsupervised clustering as these cells form distinct cluster(s) in scRNA-seq datasets and have unambiguous identities. To avoid batch effects from various datasets due to technical variabilities such as sample preparation and different sequencing platforms, we performed all our differential gene analysis on comparisons of orthologous gene expression between imGCs and mGCs within each dataset to reveal the enriched genes and their associated biological processes (two-sided Wilcoxon rank sum test; false discovery rate (FDR)-adjusted *P* < 0.05 and absolute value of log_2_-fold changes > 0.1 as significantly differentially expressed) as opposed to directly cross-comparing the expression of individual genes in imGCs of different species in different datasets^57^. We found that imGCs in humans, macaques, and pigs all showed significantly higher enrichment of the mouse imGC gene module, while their mature counterparts expressed more of the mouse mGC module genes (one-way Wilcoxon rank-sum test) (**Fig. 3a**). The imGCs in all four species predominantly enrich for genes associated with GO term groups of biological processes related to neurogenesis and neuron development (**Fig. 3b; Extended Data Fig. 4a; Supplementary Table 4**). These results indicate that imGCs in different mammalian species, including those identified using our machine learning models, have immature neuronal characteristics at the whole transcriptome level and share similar biological processes to maintain their immature properties.

**Figure 3.**
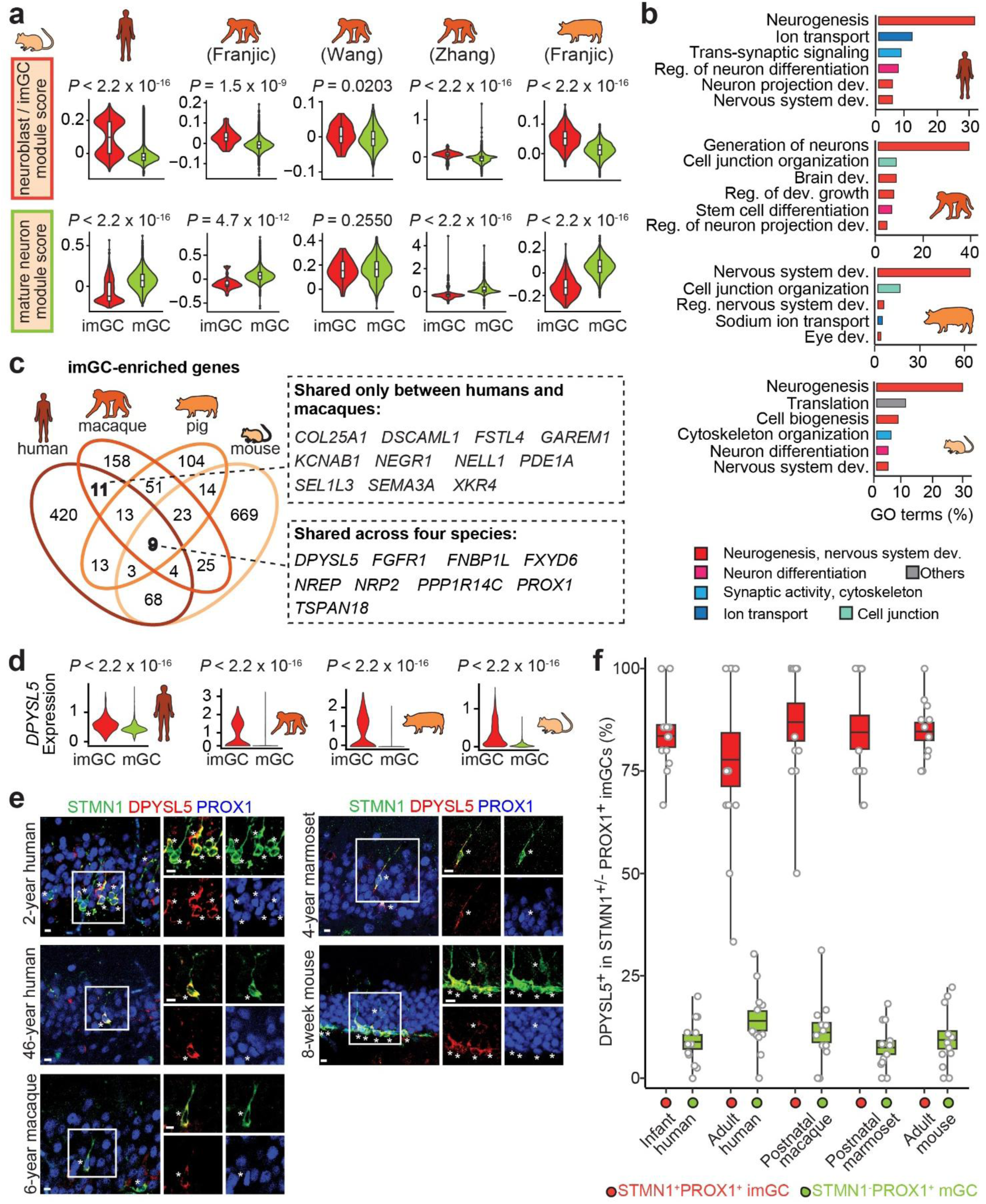
Conserved immature neuronal biological processes with divergent gene expression features in imGCs of different species. **a,** Gene module analysis of human^31^, macaque^25,27,28^, pig^28^ imGCs-enriched and mGCs-enriched genes using a published list of mouse immature progeny (neuroblast and imGC)-enriched and mGC-enriched genes^16^ (one-way Wilcoxon rank-sum test). **b,** Top GO term groups for imGC-enriched genes in different species coloured by biological processes. Dev., development; reg., regulation. **c,** Venn diagram of imGC-enriched genes in different species. Genes shared across the four species and genes shared only between humans and macaques are listed in the boxes. **d,** Violin plots of normalized expression of DPYSL5 in imGCs and mGCs of different species from published scRNA-seq datasets (one-way Wilcoxon rank-sum test). **e, f,** Sample confocal immunostaining images (**e**) and quantification (**f**) of DPYSL5 expression in STMN1^+^PROX1^+^ imGCs and STMN1^-^PROX1^+^ mGCs in the hippocampi of infant and adult humans, postnatal macaques and marmosets, and adult mice. Scale bar: 10 µm. Asterisks indicate DPYSL5^+^ cells among STMN1^+^PROX1^+^ imGCs (**e**). Dots represent data from individual sections; the centre line represents the mean, box edges show s.e.m. and whiskers extend to the maximum and minimum values (n = 4, 4, 2, 1, 4 subjects for infant humans, adult humans, macaques, marmosets, and mice, respectively) (**f**).

In contrast to such high conservation in biological processes, we observed widely divergent expression patterns among highly expressed genes enriched in imGCs across species, even between humans and macaques (37 among 541 human imGC-enriched genes (6.8%)), indicating substantial interspecies variance (**Fig. 3c**). Among the 541 human imGC-enriched genes, we identified only 9 highly expressed genes for imGCs over mGCs (1.6%) that were shared across the four species, including DPYSL5 (which encodes Dihydropyrimidinase-Like-5 protein involved in growth cone guidance and neural development; also known as CRMP5) and FXYD6 (which encodes FXYD domain containing ion transport regulator 6) (**Fig. 3c, d; Extended Data Fig. 4b**). Analysis using the public Allen Brain *in-situ* hybridization database^58^ found that 8 out of 9 genes (DPYSL5, FGFR1, FNBP1L, FXYD6, NREP, NRP2, PPP1R14C, and TSPAN18) exhibit enrichment of expression in the neurogenic subgranular zone of adult mice (**Extended Data Fig. 5a**). We further confirmed the enrichment of DPYSL5 protein expression in imGCs compared to mGCs in the hippocampi of infant and adult humans, postnatal macaques, marmosets, and mice using immunohistology (**Fig. 3d-f; Extended Data Fig. 5b**). As a comparison, we performed a similar cross-species analysis of GC-enriched genes compared to other cell types in the same datasets and observed a much larger overlap (**Fig. 3c**; **Extended Data Fig. 5c**). For example, we found 1410 shared highly expressed genes enriched in GCs between humans and macaques, representing 42.4% of the 3326 human GC-enriched genes, which contrasts with a 6.8% overlap between imGC- enriched genes in the two species (**Fig. 3c**; **Extended Data Fig. 5c**). Collectively, these results showed that imGCs in different species utilize shared immature cellular programs but exhibit divergent gene expression features, suggesting that despite gross transcriptomic similarity in biological processes across species, substantial species divergence can exist at the individual gene level^59–62^ (**Extended Data Fig. 5d**).

### Species-specific imGC features

To reveal species-specialized molecular properties of imGCs, we next examined highly expressed genes that are uniquely enriched in imGCs in each species (**Fig. 4a**). GO analysis of biological processes associated with these genes revealed ion transport and synaptic transmission as the two most dominant terms among humans, macaques, and pigs, whereas development-related terms were the most enriched in mice (**Fig. 4a; Supplementary Table 5**). Moreover, expression of risk genes of several human brain disorders was enriched in human, macaque, pig imGCs or mGCs, compared to those in mice (**Fig. 4b, c**). These results suggest that imGCs in humans, macaques, and pigs exhibit enhanced expression of genes associated with neuronal communication and plasticity, even though each species may utilize distinct genes, whereas mouse imGCs exhibit similarity in their gene enrichment patterns to a lesser degree compared to those of the other species.

**Figure 4.**
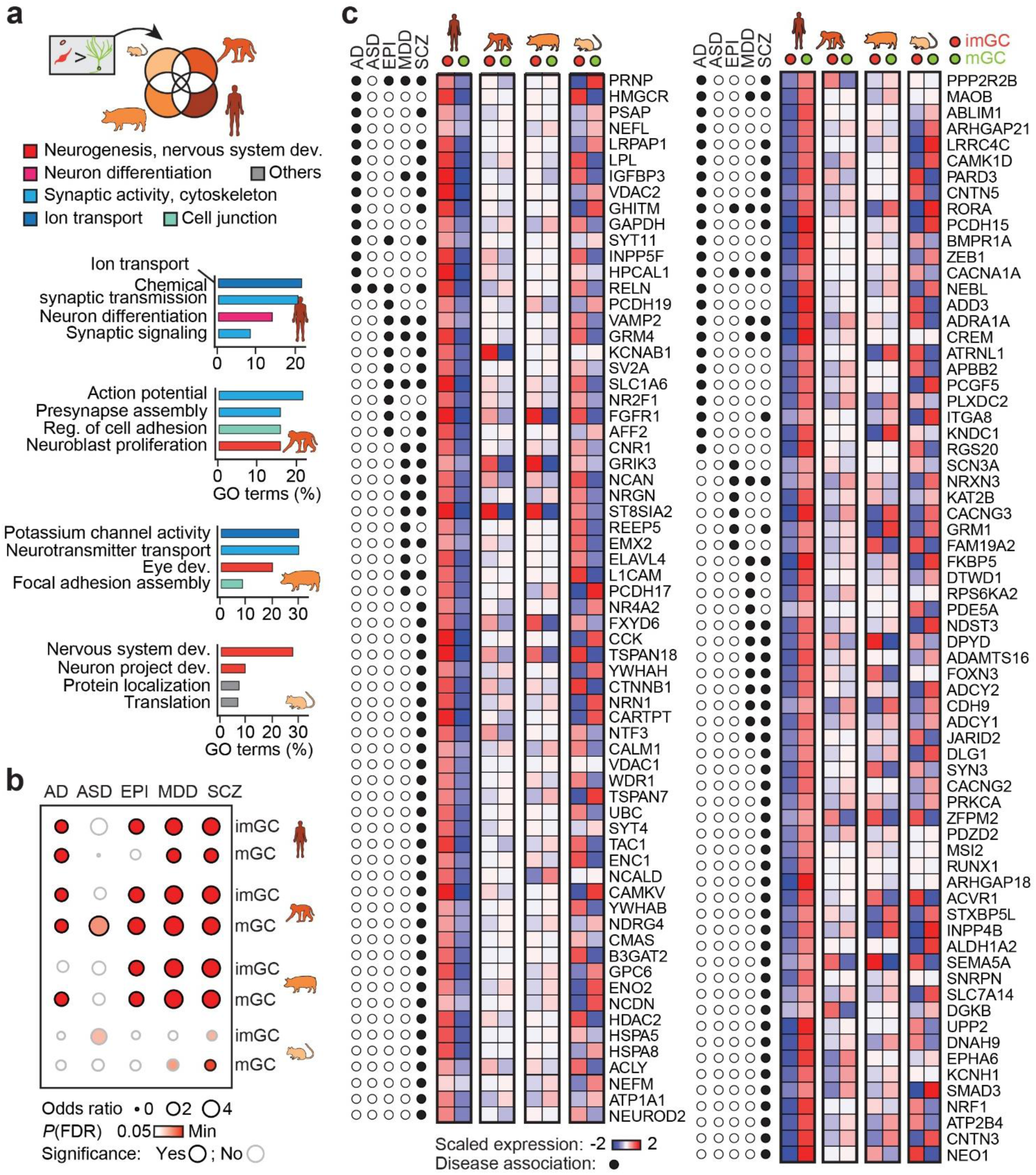
Species-specific enrichment of imGC molecular features. **a,** Schematic Venn diagram (top) and bar plots (bottom) of top GO term groups for imGC-enriched genes unique to each species coloured by biological processes. **b,** Enrichment patterns of brain disorder risk gene expression in imGCs and mGCs of different species (one-sided Fisher’s exact test, FDR-adjusted *P* < 0.05). AD, Alzheimer’s disease; ASD, autistic spectrum disorders; EPI, epilepsy; MDD, major depressive disorder; SCZ, schizophrenia. **c,** Red-blue heatmaps depict expression patterns of the risk genes of neurological or psychiatric disorders in imGCs and mGCs across four species. The colour bar represents z-scores of gene expression, scaled to range from −2 to 2.

To investigate human-enriched molecular features of imGCs in detail, we compared the expression patterns of each orthologous gene in the two major GO categories of biological processes with species-specific enrichment in imGCs and mGCs, in a non-biased manner. These two categories, “ion transport” and “synaptic transmission”, include genes that regulate various ion channels, glutamate receptors, GABA receptors, ATPases, and transmembrane proteins (**Extended Data Fig. 6**). We observed enrichment of distinct gene families in human imGCs over mGCs, such as genes encoding Na^+^/K^+^-ATPase (ATP1 gene family), shaker-related alpha 1 subunit of voltage-gated K^+^ channel (KCNA gene family), and canonical transient receptor potential cation channel (TRPC gene family) (**Extended Data Fig. 7**). Notably, most genes that encode for a distinct subtype of the ATPase family responsible for lysosomal vacuolar-type H^+^ transport (v-ATPase; ATP6 gene family) displayed imGC enrichment unique in humans but not in the other three species (**Fig. 5a**). We validated the enriched expression pattern of v-ATPase genes in imGCs over mGCs in several published scRNA-seq datasets of the human hippocampus^27–29,31,34^ (**Fig. 5a; Supplementary Table 1**). We further confirmed the enrichment of RNA molecules of two genes encoding v-ATPase in human imGCs and mouse mGCs using *in-situ* hybridization (**Fig. 5b, c**).

**Figure 5.**
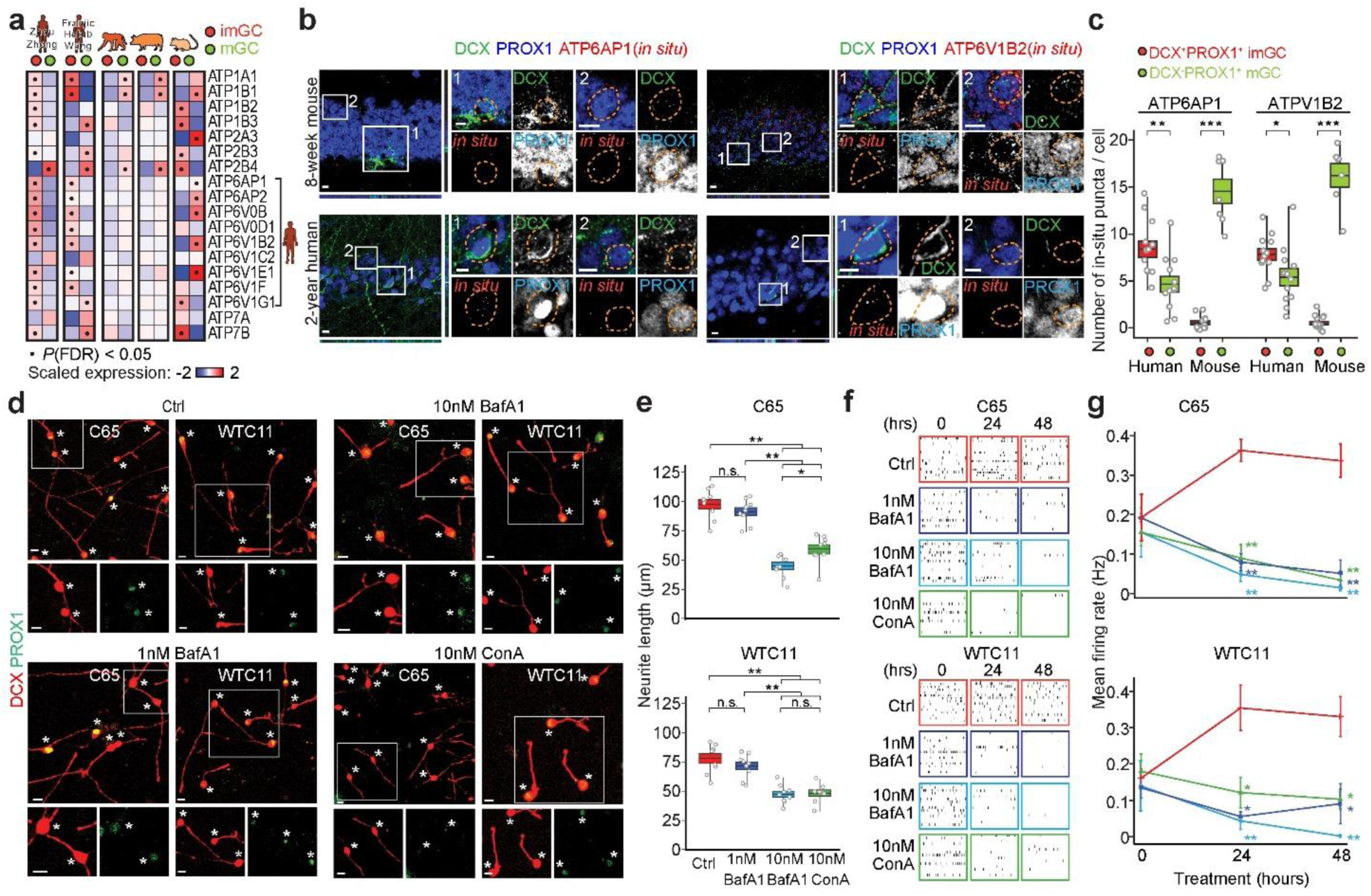
Human-enriched hippocampal immature neuron features and functional roles of a family of genes encoding lysosomal vacuolar-type H^+^-transporting ATPases (v-ATPases) in their development. **a,** Expression patterns of genes encoding ATPases in imGCs and mGCs in different species (black dots indicate significant difference with FDR-adjusted *P* < 0.05 using two-sided Wilcoxon rank sum test). Several previously published scRNA-seq datasets of the human hippocampus (left two columns) were used to ensure consistency across studies in gene expression enrichment patterns^27–29,31,34^. The colour bar represents z-scores of gene expression, scaled to range from −2 to 2. **b, c,** Sample confocal *in-situ* hybridization and immunostaining images (**b**) and quantification (**c**) of expression of ATP6AP1 and ATP6V1B2, two subunits of v-ATPases, in human and mouse GCs. Orange circles in image sets #1 indicate the presence and absence of ATP6AP1 and ATP6V1B2 *in situ* puncta signals in human and mouse imGCs (DCX^+^PROX1^+^), respectively (**b**). Orange circles in image sets #2 indicate the absence and presence of ATP6AP1 and ATP6V1B2 *in situ* puncta signals in human and mouse mGCs (DCX^-^PROX1^+^), respectively (**b**). Dots represent data from individual fields of view; the centre line represents the mean, box edges show s.e.m. and whiskers extend to the maximum and minimum values (**c**) (n = 2 subjects each for humans and mice; Student’s t-test, * *P* < 0.05, ** *P* < 0.005, *** *P* < 0.0005). **d, e,** Sample confocal immunostaining images (**d**) and quantification (**e**) of neurite length of DCX^+^PROX1^+^ imGCs derived from two independent human iPSC lines. *In vitro* culture was treated with Bafilomycin A1 (BafA1) and Concanamycin A (ConA), two specific blockers against v-ATPases for 24 hours (hrs). See **Extended Data Fig. 8a**. Asterisks indicate DCX^+^PROX1^+^ imGCs (**d**). Scale bar, 10 µm (**d**). Box colours match treatment conditions. Dots represent data from individual images; the centre line represents the mean, box edges show s.e.m. and whiskers extend to the maximum and minimum values (n = 3 cultures per condition). One-way ANOVA with post-hoc Tukey HSD test, * *P* < 0.05, ** *P* < 0.005 (**e**). **f, g,** Sample raster plots (**f**) and quantification (**g**) of neuron firing rate of hippocampal imGCs using multi-electrode array assay with or without drug treatments. See **Extended Data Fig. 8a**. Values represent mean ±s.e.m. (n = 3 per condition; one-way ANOVA with post-hoc Tukey HSD test was used to compare among treatment conditions, * *P* < 0.05 or ** *P* < 0.005 when compared to the DMSO control group (Ctrl), and all other pair-wise comparisons were not significant).

### Neuronal development regulated by human imGC-enriched lysosomal v-ATPases

We next explored functional roles of human-enriched expression of v-ATPases in imGCs (**Extended Data Fig. 8a**). We developed *in vitro* culture models of DCX^+^PROX1^+^ human hippocampal imGCs derived from two human iPSC lines of different genetic backgrounds and confirmed their imGC characteristics using immunohistology (**Extended Data Fig. 8a-e**). To examine the collective influence of the many gene family members enriched in human imGCs, we used two specific pharmacological blockers of v-ATPases^63^, Bafilomycin A1 (BafA1)^64^ and Concanamycin A (ConA)^65^. We found that these two blockers did not have an effect on imGC identity or cell survival at 1 nM or 10 nM (**Extended Data Fig. 8b-e**). We further examined the acute effects of these v-ATPase blockers on imGC neuronal development by measuring neurite length and neuronal firing rate after a short duration of treatment (**Extended Data Fig. 8a**). Addition of 10 nM of either BafA1 or ConA leads to a significant decrease in neurite growth whereas 1 nM has no effect at 24 hours after treatment (**Fig. 5d, e**). Moreover, imGC firing rate measured by multi-electrode array assay showed a significant decrease upon BafA1 or ConA treatment at 1 nM and 10 nM after 24 or 48 hours (**Fig. 5f, g**). These results showed that lysosomal v-ATPases, which exhibit human-specific enriched expression in imGCs, play functional roles in human imGC neuronal development.

## Discussion

Here we made a surprising discovery of substantial differences in the highly expressed genes enriched in imGCs relative to mGCs in the hippocampus among humans, macaques, pigs, and mice and identified human-specialized features that have functional roles in human neuronal development. Interestingly, despite interspecies variance in highly expressed genes with imGC- enriched expression, the associated GO biological processes of these genes converged to common biological processes regulating neurogenesis, neuron development, and plasticity, suggesting that imGCs utilize species-unique molecular features to achieve similar cellular maintenance of immature neuronal characteristics. Our findings suggest a new concept in neuroscience, namely, biological processes are conserved across different species, while the regulation of individual genes within these processes can be vastly divergent, potentially allowing for phenotypical and functional adaptations (**Extended Data Fig. 5d**). A similar concept has been suggested by several previous studies in the evolutionary biology field, where convergently evolved phenotypic traits have occurred through genotypically distinct regulatory changes, for example, in tooth gain in two different types of fish^59^, viviparity in eight types of vertebrate species^60^, chemosensory tissue development in six *Drosophila* species^61^, phenotypic traits adapted to aquatic environment among killer whales, walruses and manatees^62^, and iPSC proliferation between humans and chimpanzees^66^. However, such a concept of “recruitment of divergent genes in different species resulting in regulation of a convergent biological process” is underappreciated in the neuroscience field and is conceptually different from most findings from recent studies of the developing cortex, in which a single genetic variant can be responsible for cross-species cellular innovations of new neuron regulation^39^.

Cross-species transfer learning approaches for cell typing in scRNA-seq analyses have been commonly used in the field, which assume general similarities of gene expression in cell types across species and is very effective in distinguishing broad cell types^67,68^. However, this approach can pose challenges when attempting to discern differences between cellular subtypes or cellular states that exhibit subtle variations^31,68–71^. Given the limited capability of the traditional unsupervised clustering or transfer learning methods in separating cell types with fine molecular differences, as shown by inconsistent results in identifying macaque imGCs^30^, caution needs to be exercised in assuming molecular similarity between species or applying molecular signatures from one species to others for cell type annotations. Here, we trained a machine learning-based model with high-confidence immature neuron prototypes from the young macaque hippocampus to identify macaque imGCs in scRNA-seq datasets, which provides a transcriptome-wide definition of macaque imGCs *in silico* using a combinatorial weighted gene panel, rather than arbitrarily picked marker gene lists (**Fig. 1a**). Using such approaches to identify imGCs in macaques and humans^31^ enabled us to identify cells exhibiting consistent immature, neuronal, regional (dentate granule) characteristics despite potential marker gene negativity that could result from species-specific gene expression variations, or low depth or stochastic detection by scRNA-seq. Many conventional markers identified in mice, such as DCX and STMN1, exhibited a tendency toward enrichment in imGCs but did not achieve statistical significance among the imGC-enriched genes shared across the four species studied. For example, DCX was significantly enriched in imGCs of mice, pigs and macaques, but not in humans. These results may be attributed to either low or stochastic mRNA expression levels, or to limitations in detection by scRNA-seq technologies^31,38^ (**Fig. 3c**). In addition, we noticed a considerable degree of variability among individuals in the average proportion of imGCs among GCs in both humans^31^ and macaques (**Fig. 2b**). We cannot rule out the effect from the low number of specimens sequenced and their heterogeneous backgrounds. Owing to inherent limitations in sequencing depth associated with scRNA-seq technology, our analysis focuses only on highly expressed genes, therefore, we do not know whether genes enriched in imGCs over mGCs but with lower expression and not captured by scRNA-seq exhibit cross-species similarity or differences. We also did not consider species-unique genomic features or isoform usage. In the future, sequencing more samples more deeply and integrating multiple data modalities – such as co-detection of transcripts and chromatin accessibility, pre-labelling and/or enrichment with antibodies, nucleotide analogues, or barcodes, and long-reads sequencing to detect differential exon/isoform usage – could mitigate variability due to sampling and allow for further investigation of imGC similarities and differences, such as males versus females, and prenatal versus postnatal across species.

Our work has revealed many human-specific imGC features in gene expression and developmental dynamics. We identified unique sets of differentially and highly expressed genes in imGCs over mGCs in humans, including genes encoding an ATPase subfamily for lysosomal H^+^ transport (**Fig. 5a**). We validated the collective functional roles of the lysosomal H^+^-transporting v-ATPase, encoded by genes in the ATP6 family, in human imGC development and function *in vitro* via specific pharmacological inhibition using human iPSC-derived hippocampal imGCs (**Extended Data Fig. 8a**). Since this *in vitro* model does not fully recapitulate the process of adult neurogenesis *in vivo*, the roles of these genes in regulating imGC development during adult human neurogenesis remains to be determined. Several subunits of the v-ATPase have been shown to regulate neural development and have neurological disease implications, such as those encoded by ATP6AP1^66^, ATP6AP2^66,72–76^, ATP6V1E1^77,78^, among others^79–82^. Notably, cell growth mediated by mTORC1 signalling was perturbed upon ATP6AP1 knockout in human cells but was significantly less in chimpanzees, indicating human-specific sensitivity^66^. It is important to note that among the genes differentially expressed in imGCs over mGCs specifically in humans, some may also exhibit a lower level of expression in human mGCs or mouse imGCs. Therefore, they may potentially play a role in these cells as well. Recent studies have identified multiple human brain regions with postnatal development of immature neurons, such as in the amygdala or entorhinal cortex^83,84^, indicating special temporal dynamics in human neuronal development. These studies underscore the significance of conducting cross-species comparative analysis to identify molecular signatures that are differentially enriched in humans. Together, our findings emphasize the utility of human cell-based models, including cultures derived from human stem cells and surgically resected human samples, in examining the molecular characteristics, functionalities, and regulatory mechanisms of immature neurons in humans^37,39,57^. It is also crucial to use human cell-based models to validate therapeutic strategies aimed at regulating adult hippocampal neurogenesis in patients.

Taken together, our study presents the comparative molecular landscapes of immature neurons in the mammalian hippocampus across various species and highlights substantial interspecies variance in highly expressed genes enriched in imGCs over mGCs, but conserved and convergent cellular programs regulating neuronal development. Our results also highlight the importance of conducting independent molecular and functional analyses for adult neurogenesis in different species and provide the foundation for future molecular analysis and targeting of imGCs to facilitate our understanding of primate and human neuronal development.

## Methods

### Tissue specimens

De-identified human tissue specimens were collected and processed under protocols approved by the Institutional Review Boards (IRB) of the University of Pennsylvania, the Children’s Hospital of Philadelphia, and Center for Excellence in Brain Science and Intelligence Technology at the Chinese Academy of Sciences. A total of 10 human hippocampal specimens were used for immunohistological analyses (**Supplementary Table 6**). Postnatal postmortem hippocampal specimens were collected from tissue banks at the Children’s Hospital of Philadelphia, the Johns Hopkins University Pathology Archive, the Lieber Institute for Brain Development, the NIH NeuroBioBank at the University of Pittsburgh Brain Tissue Donation Program, the University of Maryland Brain and Tissue Bank, and the China Human Brain Bank Consortium. Informed consent for each specimen was obtained by its corresponding institution prior to tissue collection.

For mouse immunohistological analysis, adult (8 weeks old), wild-type, male and female, C57BL/6 mice were used. No obvious sex-dependent phenotype was observed in any of the experiments. Animals were housed in a 12-hour dark/ light cycle with food and water ad libitum. For immunohistological analysis of the non-human primate hippocampus, post-mortem tissue from macaques (*Macaca fascicularis*; also known as cynomolgus monkeys) and marmosets (*Callithrix jacchus*) were used. Animal procedures were performed in accordance with protocols approved by the Institutional Animal Care and Use Committee of the University of Pennsylvania and the Biomedical Research Ethics Committee of Center for Excellence in Brain Science and Intelligence Technology at the Chinese Academy of Sciences.

### Dataset pre-processing, integration, and clustering

Single-cell or single-nucleus RNA sequencing (scRNA-seq) datasets were processed as previously described with minor modifications^31^. Unique Molecular Identifiers (UMI) count matrices from published datasets^16,25–29,31,34,54^ were retrieved from the respective repositories and processed independently using the same criteria (summarized in **Supplementary Table 1**). Count matrices were loaded into the R (v4.1.3; https://www.r-project.org/) package ‘Seurat’ (v4.3.0)^51^. To mitigate the impact of batch effects from technical variabilities across different datasets (such as varying sequencing platforms, sequencing depth), which could hinder the performance of our machine learning-based classification of macaque imGCs, we ensured that each specimen of the macaque datasets exhibited comparable sequencing characteristics by performing random down-sampling (implemented in the “rbinom” function in R) on expression matrices with significantly higher average number of genes and reads per cell (> 2,000 genes/cell and > 4,000 reads/cell) to achieve a similar sequencing depth as the macaque hippocampal dataset with the lowest number of genes and reads per cell (∼1,470 genes/cell and ∼2,700 reads/cell) prior to quality control and downstream processing. This process was repeated 10 times to ensure robustness and consistency.

For each specimen, genes expressed in < 10 nuclei were discarded. Nuclei with < 400 or > 5,000 genes were discarded; nuclei with > 5% UMIs mapped to mitochondrial genes were discarded. For normalization, UMI counts for all nuclei were scaled by library size (total UMI counts), multiplied by 10,000 and transformed to a log scale. Highly variable genes were identified using the function ‘FindVariableFeatures’ in Seurat. The top principal components (PCs), determined by the ‘PCElbowPlot’ function, were selected for dimensionality reduction, clustering, and visualization with Uniform Manifold Approximation and Projection (UMAP). Marker genes for each cluster were identified with a Wilcoxon rank sum test implemented in the ‘FindAllMarkers’ function with the following criteria: false discovery rate (FDR)-adjusted *P*-value < 0.01, log_2_-fold change ≥ 0.5, and genes detected in < 25% of the cells within its corresponding cluster were excluded. For example, PROX1 was used to determine the cell identities of excitatory dentate granule cells in hippocampal scRNA-seq datasets.

Datasets containing multiple specimens were normalized within each study to remove sequencing variation (implemented in ‘sctransform’ function in Seurat^85^) prior to integration using a canonical correlation analysis (CCA) in Seurat^51^. For integrated datasets, the top 2,000 highly variable genes and the first 30 principal components were used for cell alignment before clustering and UMAP visualization.

### Macaque imGC signature extraction and prototype-based scoring using machine learning

To ensure a precise transcriptome-wide characterization of macaque imGCs, considering substantial species differences and the potential technical variability in different published datasets, we implemented a supervised learning approach^31^. This approach builds a model to learn comprehensive gene features from highly confident macaque imGCs (serving as cell prototypes for model training), which we then used to quantitatively assess the similarity of each cell in query (test) scRNA-seq datasets. A multinomial machine learning method using a L2-norm regularized logistic regression model^86^ (implemented in the ‘LogisticRegression’ function in ‘scikit-learn’^87^ in Python v3.7) was applied with modifications^31,86^. The sensitivity and specificity of this approach were validated in separate models established to identify human and mouse imGCs^31^.

To construct a prototype-based scoring model for the precise characterization of macaque imGCs, we selected cell-type prototypes from an unsupervised-clustered scRNA-seq dataset of the young (4-6 years, neurogenic) macaque hippocampi^25^ as the training dataset. The prototypes consist of imGCs and all major non-neuronal cell types, including astrocytes (Astro), oligodendrocyte precursor cells (OPC), mature oligodendrocytes (mOli), and microglia. Since DCX transcripts are not solely present in the GC cluster, we selected the imGC prototypes based on their defining gene expression features, DCX^+^CALB1^-^PROX1^+^, within the GC cluster to ensure prototype purity and representativeness. Other cell-type prototypes were chosen by their defining features within their respective clusters, including AQP4^+^ cells from the astrocyte cluster, PDGFRA^+^ cells from the OPC cluster, MOBP^+^ cells from the mature oligodendrocyte cluster, and CX3CR1^+^ cells from the microglia cluster. To further ensure representativeness of cell-type prototypes, two negative selection criteria were employed. Firstly, cells expressing common markers of the other prototypical cell types were excluded from prototypes. For example, imGC prototypes were required to lack expression of markers of astrocytes (SLC1A2, AQP4), OPCs (PDGFRA), oligodendrocyte lineage cells (OLIG2, CNP, MBP, MOBP), and microglia (CX3CR1, PTPRC). Secondly, cells were excluded from all prototypes if they expressed defining markers of other known cell types in the hippocampal dataset, including GABAergic interneurons (GAD1, GAD2), *cornu ammonis* 1 excitatory neurons (SATB2), ependymal cells (FOXJ1), endothelial cells (FLT1), and blood cells (HBA1).

The cell scoring model was trained on a log-transformed, max-normalized count matrix of the prototype cells with all genes retained, followed by a gene ranking procedure^86,88^ to refine for highly variable cell-type specific markers. An optimal regularization parameter of 0.007 for the logistic regression model was determined by plotting the regularization strength against the classifier accuracy, looking for the most stringent value of regularization with a maximal accuracy rate (∼99.5%). An in-built cross-validation procedure was applied to the training set to estimate the average accuracy of the model (implemented in the ‘LogisticRegressionCV’ function in ‘scikit-learn’) with the following parameters, training set:validation set = 85%:15%, and training set randomly split for 35 iterations (using a stratified k-fold cross-validation approach). The trained model employs a list of positively-weighted and negatively-weighted coefficients to prioritize genes based on their predictive capacity for each cell type category. A key aspect of this model is its reliance on a combinatorial gene panel with different weights, rather than a limited set of arbitrarily selected markers, to characterize the transcriptomic profile of macaque imGCs. This approach ensures a balanced representation of immature, neuronal, and regional (dentate gyrus) features, providing a more comprehensive understanding of the cellular characteristics defining imGCs (**Supplementary Table 2**).

All accessible published scRNA-seq datasets of the macaque hippocampus and one from the young adult “non-neurogenic” neocortex (serving as a negative control) were retrieved from public repositories and individually prepared using Seurat^51^ as query (test) datasets (**Supplementary Table 1**). Utilizing the trained model, we evaluated the probabilistic similarity (*p*) of each individual cell from the test datasets (log-transformed and max-normalized count matrices) to each prototype. This assessment was conducted without any prior knowledge of cell clustering information. The predicted probabilities for each cell in the test datasets, ranging from 0 to 1, were determined using the ‘softmax’ function (implemented in the ‘predict_proba’ function in ‘scikit-learn’)^86^. Similarity scores were projected onto their corresponding UMAP clustering plots, allowing a visual representation of the cells’ similarity to prototype imGCs (**Fig. 2a**). To classify a cell as a macaque imGC, we empirically established a conservative cutoff of *p_(imGC)_* ≥ 0.85, of which the effectiveness and specificity was validated through its ability to identify human and mouse imGCs^31^.

### Comparison of imGC and mGC features by gene module analysis and differential gene expression analysis

To avoid batch effects from datasets due to technical variabilities (such as different sequencing platforms), we performed all gene comparisons between imGCs and mGCs within individual datasets rather than directly cross-comparing the expression of individual genes in imGCs of different species. Among the four species, human imGCs (identified in our previous study^31^) and macaque^25,27,28^ imGCs were identified using their respective machine learning scoring models; for pig^28^ and mouse^16^ imGCs, we used annotations based on unsupervised clustering as they form distinct cluster(s) in scRNA-seq datasets and have unambiguous identities.

To evaluate the immature characteristics of imGCs and mGCs in different species, we performed a gene module analysis (implemented in the ‘AddModuleScore’ function in Seurat^51^) using a well-acknowledged list of mouse immature progeny (neuroblasts and imGCs) and mGC signatures^16^.

Differentially expressed genes (DEGs) were identified using a two-sided Wilcoxon rank sum test (implemented in the “FindMarkers” function in Seurat), using the ‘RNA slot’ of the Seurat object. To examine the differential gene features between imGCs and mGCs, we included all imGCs and mGCs and directly contrasted them within each integrated dataset. To validate the robustness of scRNA-seq analysis of the same datasets, we compared all GCs (imGCs and mGCs) to all other cell types within each dataset (**Extended Data Fig. 5c**). To ensure a fair statistical comparison for all analyses, “max.cells.per.ident” in the “FindMarkers” function was determined by the cell number of the group with fewer cells. Genes with FDR-adjusted *P*-value < 0.05 and fold change (log_2_ scale, absolute value) > 0.1 were considered significantly differentially expressed. For DEG comparison across species, only genes with orthologs to humans in all four species (identified using the ‘babelgene’ (v22.9; https://igordot.github.io/babelgene/) package in R) were included. To minimize biases introduced by variations in sequencing platforms across studies, we utilized a dataset from a single macaque study^25^ that has a significantly larger number of specimens compared to other available datasets, as well as young and old age groups, for DEG analysis of macaque imGCs and mGCs unless otherwise specified.

### Gene ontology and disease risk gene enrichment analyses

Gene ontology (GO) networks of biological processes were analysed using the ClueGO (v.2.5.5) plug-in^89^ in Cytoscape^90^ (v3.7.2) with the following settings: ‘GO Biological process’ of each corresponding species was selected (accessed on March 21, 2023); running the default one-sided hypergeometric test, only pathways with FDR-adjusted *P*-value < 0.05 were displayed; ‘GO fusion’ option was enabled; ‘% Associated Genes’ are no less than 3%. Genes identified in the machine learning model or from differential expression analyses were selected as the significantly regulated genes and used as input. For groups with more than 200 significantly regulated genes, a minimum of 7 genes per cluster were used; for groups with less than 100 significantly regulated genes, a minimum of 2 genes per cluster were used; and for all other groups, a minimum of 3 genes per cluster were used.

Lists of genes associated with GO term groups, “ion transport” (GO_0098660) and “synaptic transmission” (GO_0035249), were obtained from http://www.informatics.jax.org/ on January 2, 2023 (**Extended Data Figs. 6, 7**).

To map risk genes for brain disorders, we analysed the enrichment and aggregated expression (**Fig. 4b, c**) of the significant DEGs in each category with disease annotations collected from the Phenopedia database^91^ (accessed on March 25, 2021). For disease risk gene enrichment analysis, we calculated the odds ratios and the enrichment p-values, which were determined by a one-tailed Fisher’s exact test (implemented in the ‘fisher.test’ function in R) and corrected by controlling for the FDR for multiple comparisons.

### Human hippocampal neuron *in vitro* culture analysis

Hippocampal dentate granule cell-like neurons were generated from human pluripotent stem cells (iPSCs)^92^ (**Extended Data Fig. 8a**). Human iPSC lines used in the current study were derived from healthy donors as previously described and were either obtained from commercial sources or previously generated and fully characterized^93–96^. C65 human iPSCs were generated from fibroblasts of healthy adult subjects^94,96^; WTC11 human iPSCs were obtained from Coriell (GM25256). All procedures were performed in accordance with IRB and ISCRO protocols approved by the Institutional Committees. Human iPSCs were maintained in mTeSR Plus (STEMCELL Technologies) on Matrigel (Corning)-coated plates and dissociated by TrypLE (Gibco) at Day 0. Approximately 3×10^6^ cells were aggregated to form ∼300 embryoid bodies (EBs) by Aggrewell800 (STEMCELL Technologies). The following day, EBs were resuspended and transferred to 6-well plates rotating at 110 rpm and patterned into an ectoderm fate in neural induction media for 5 days^97,98^. From Day 6 to Day 13, EBs were patterned to a dentate granule cell fate in hippocampal patterning media^97,98^ rotating at 110 rpm with half of the medium changed daily. From Day 14, EBs were cultured in neural progenitor media^97,98^ rotating at 110 rpm with half of the medium changed daily. On Day 16, EBs were dissociated into single cells by TrypLE and plated on Matrigel-coated plates at a density of 2×10^6^ cells per well on 6-well plates for neural progenitor expansion. Medium was changed every two days. After one week, neural progenitors were dissociated and seeded at a density of 2×10^5^ cells per well on 24-well plates with Poly-D-Lysine and laminin-coated coverslips (Sarstedt Inc.) in neuron culture media^97,98^ for neuron differentiation. For Bafilomycin A1 (BafA1) and Concanamycin A (ConA) treatment, neurons differentiated for 1 week were treated with BafA1, ConA or DMSO for 24 hours or 48 hours. The Axion Maestro system was used for multi-electrode array (MEA) recording. Neural progenitors were seeded at a density of 5×10^4^ cells per well into 24- well MEA plates and differentiated for 2 weeks. Neuronal activities were recorded before and after treatment.

### Immunostaining and confocal microscopy

Brain tissue sections were pre-treated and immunohistology was performed as previously described with minor modifications^31^. In brief, brain tissue blocks were fixed with 4% paraformaldehyde (PFA) at 4 °C for 24-48 hours, and cryoprotected with 30% sucrose (wt/vol). 40-µm-thick sections were cut on a frozen sliding microtome (Leica, SM2010R) as previously described^99^. The sections then underwent antigen retrieval prior to antibody application by being incubated in 1×target-retrieval solution (DAKO) at 95 °C for 12.5 minutes, followed by 15 minutes of cooling to room temperature. Coverslips of *in vitro* culture of iPSC-derived hippocampal imGCs were fixed with PFA for 30 min at room temperature prior to antibody incubation^93,99^. Antibodies were diluted in Tris buffered saline (TBS) with 0.1% Triton X-100, 5% (vol/vol) donkey serum (Millipore, S30), and sodium azide (Sigma, S2002, 1:100). Sections were incubated with primary antibodies at 4 °C for two days. The following primary antibodies were applied: cleaved caspase 3 (rabbit, Cell Signaling Technology, 9661s, 1:250), DPYSL5 (CRMP5, rabbit, Novus Biologicals, NBP1-33419, 1:300), doublecortin (DCX, guinea pig, Millipore, AB2253, 1:500), doublecortin (DCX, rabbit, Cell Signaling Technology, 4604s, 1:500), OP18 (STMN1, mouse, Santa Cruz Biotechnology, sc48362, 1:250), PROX1 (mouse, Millipore, MAB5654, 1:500), PROX1 (goat, R&D Systems, AF2727, 1:500), and STMN1 (goat, GeneTex, GTX89411, 1:500). The cyanine (Cy)-conjugated secondary antibodies raised in donkey (Jackson ImmunoResearch; 1:300), including Cy2 anti-goat (705-225- 147), mouse (715-225-151), rabbit (711-225-152), Cy3 anti-goat (705-165-147), mouse (715-165-151), rabbit (711-165-152), Cy5 anti-goat (705-175-147), mouse (715-175-151), and rabbit (711- 175-152), were incubated at room temperature for 2 hours along with DAPI (Roche, 10236276001, 1:1000). After washing with TBS, adult human brain sections were incubated with 1×TrueBlack (Biotium, 23007; diluted 1:20 in 70% ethanol) for 1 min to block the autofluorescent lipofuscin and blood components. After washing with phosphate buffered saline (PBS), stained sections were mounted and imaged as Z-stacks on a Zeiss LSM 800 confocal microscope (Carl Zeiss) using a 20× or a 40×objective with Zen 2 software (Carl Zeiss).

### *In-situ* hybridization with immunostaining

*In-situ* hybridization was performed using the RNAscope™ Multiplex Fluorescent Reagent kit v2 according to manufacturer specifications (Advanced Cell Diagnostics (ACD), 323100). Briefly, 40- µm-thick, PFA-fixed and cryopreserved brain tissue sections were mounted on Superfrost^TM^ Plus slides and baked at 60 °C for 30 minutes, followed by 15-minute post-fixation with PFA and dehydration by ethanol gradient (50%, 70%, and 100%). Tissue was then treated with hydrogen peroxide for 10 minutes for blocking of endogenous peroxidases, followed by hybridization target retrieval by incubating tissue in slightly boiling 1×RNAscope™ Target Retrieval Reagents (ACD, 322000) for 10 minutes. Tissue was treated with RNAscope™ Protease Plus (ACD, 322331) for 30 minutes prior to *in-situ* hybridization and amplification. Tissue was probed with ATP6AP1 (ACD, 1190161-C3) or ATP6V1B2 (ACD, 1584111-C1, 804291) probes and developed with TSA Vivid Fluorophore Kit 650 (ACD, PG-323273, 1:1,500). Some of the *in-situ* hybridization experiments were performed using HCR™ RNA-FISH technology^100^ following their protocol for fresh/fixed frozen tissue sections (https://www.molecularinstruments.com/hcr-rnafish-protocols). Probes for HCR™ RNA-FISH were ordered as oligoarray complex pools (Tsingke Biotech: [ATP6AP1 human]: SHP24051100014; [ATP6AP1 mouse]: SHP24051100015).

Following *in-situ* hybridization, sections were washed in PBS with 0.1% TritonX-100 (PBST), and subsequently blocked with 10% donkey serum-PBST for 1 hour at room temperature. Slides were then incubated with antibodies against doublecortin (DCX, rabbit, Cell Signaling Technology, 4604s, 1:500) and PROX1 (goat, R&D Systems, AF2727, 1:500) in 1% donkey serum PBST overnight at 4 °C, washed and then incubated with fluorescently conjugated secondary antibodies (donkey anti-goat AlexaFluor 488 [Invitrogen, A32814] and donkey anti-rabbit AlexaFluor 555 [Invitrogen, A31572], 1:500) for 2 hours at room temperature. Slides were washed with PBS and coverslipped with DAPI. Sections were imaged by confocal as described above.

### Image processing and data analysis

All confocal images were blindly acquired among different specimens under the same laser power and gain, and analysed as Z-stacked images using Imaris software (v7.2 and v9.0; BitPlane) as previously described^101,102^. The Spots module in Imaris was used to digitize cell-nucleus locations in 3D space and to code cell type classifications according to distinct morphological and molecular markers. A minimum of three randomly chosen areas of equal dimensions within each image was quantitated. The sum of quantifications of these areas per image was considered as one data point. No statistical methods were used to predetermine sample size.

*In-situ* puncta signals were blindly quantified as previously described with minor modifications^103^. To ensure consistency, all images were acquired using identical confocal settings and analyzed on a single frame basis (not z-stacked) with ImageJ (NIH). Prior to puncta counting, a cytoplasmic mask was created, and the number of puncta was extracted with the automatic ‘analyze particles’ algorithm. Each data point plotted on the graphs represents the average number of puncta observed within one cell.

### Quantification and statistical analyses

The studies were blinded during data collection and quantification. Data in figure panels reflect several independent experiments performed on different days. No data were excluded. An estimate of variation within each group of data is indicated using standard error of the mean (s.e.m.). All data are shown as mean ±s.e.m. All statistical analyses are indicated in the text or Figure legends, performed with the R language for Statistical Computing.

## Supporting information

Zipped Supplementary Tables

## Data availability

All single-cell or single-nucleus RNA sequencing data were previously published (summarized in **Supplementary Table 1**). Additional information required for reanalysing the data presented in this study can be obtained from the corresponding authors upon request.

## Code availability

Scripts for bioinformatic analyses used in this study are available at https://github.com/zhoujoeyyi/imGC_species/.

## Acknowledgements

We thank members of the Song and Ming laboratories, Zhou laboratory, and Shujia Zhu for discussion; Kimberley M. Christian for comments; Zhiheng Xu, Xiaodong Teng (ACD, Biotechne), Liqin Gu and Chao Chen (from S. Xu lab), and Lianshun Xie for help on *in situ* hybridization experiments; Brian Temsamrit, Emma LaNoce, Alan Garcia-Epelboim, Giana Alepa, and Yisha Lu for technical support; Angelina Angelucci for lab coordination; Wenqin Luo, Qinxue Wu, and Yingshi Chen for providing reagents. Some schematic illustrations were modified using images from www.biorender.com. This work was supported by grants from the National Institutes of Health (R35NS097370 and R35NS137480 to G.-l.M., R35NS116843 and RF1AG079557 to H.S., and R01NS127913 to Y.S.), The Lieber Institute for Brain Development (to J.E.K., T.M.H., and D.R.W.), Dr. Miriam and Sheldon G. Adelson Medical Research Foundation (to G.-l.M.), the National Key Research and Development Program of China (2024YFA1803400 to Y.Z.), Shanghai Science and Technology Development Funds (24QA2710400 to Y.Z.), and Shanghai Pujiang Program (23PJ1414500 to Y.Z.).

## Author contributions

Y. Zhou led the project and contributed to all aspects. Y.S., T.G., X.M., and L.W. contributed to computational data analyses. Q.Y. and Y.H. developed the *in vitro* human hippocampal neuron culture system and Q.Y. contributed to *in vitro* culture experiments. J.L., Y. Zhong, M.J., X.L., L.Y., C. Li, S.X., and J.H. contributed to histology analysis. B.C.K., A.N.V., I.H., S.K.K., J.E.K., T.M.H., D.W.N., and D.R.W. provided human hippocampal specimens. C. Liu provided marmoset specimens. Y. Zhong, N.Y., Z.L., Z.S., and C. Li provided macaque specimens. Y. Zhou, Y.S., G- l.M., and H.S. conceived the project and wrote the manuscript with inputs from all authors.

## Competing interests

The authors declare no competing interests.

## Extended Data Figures

**Extended Data Fig. 1.**
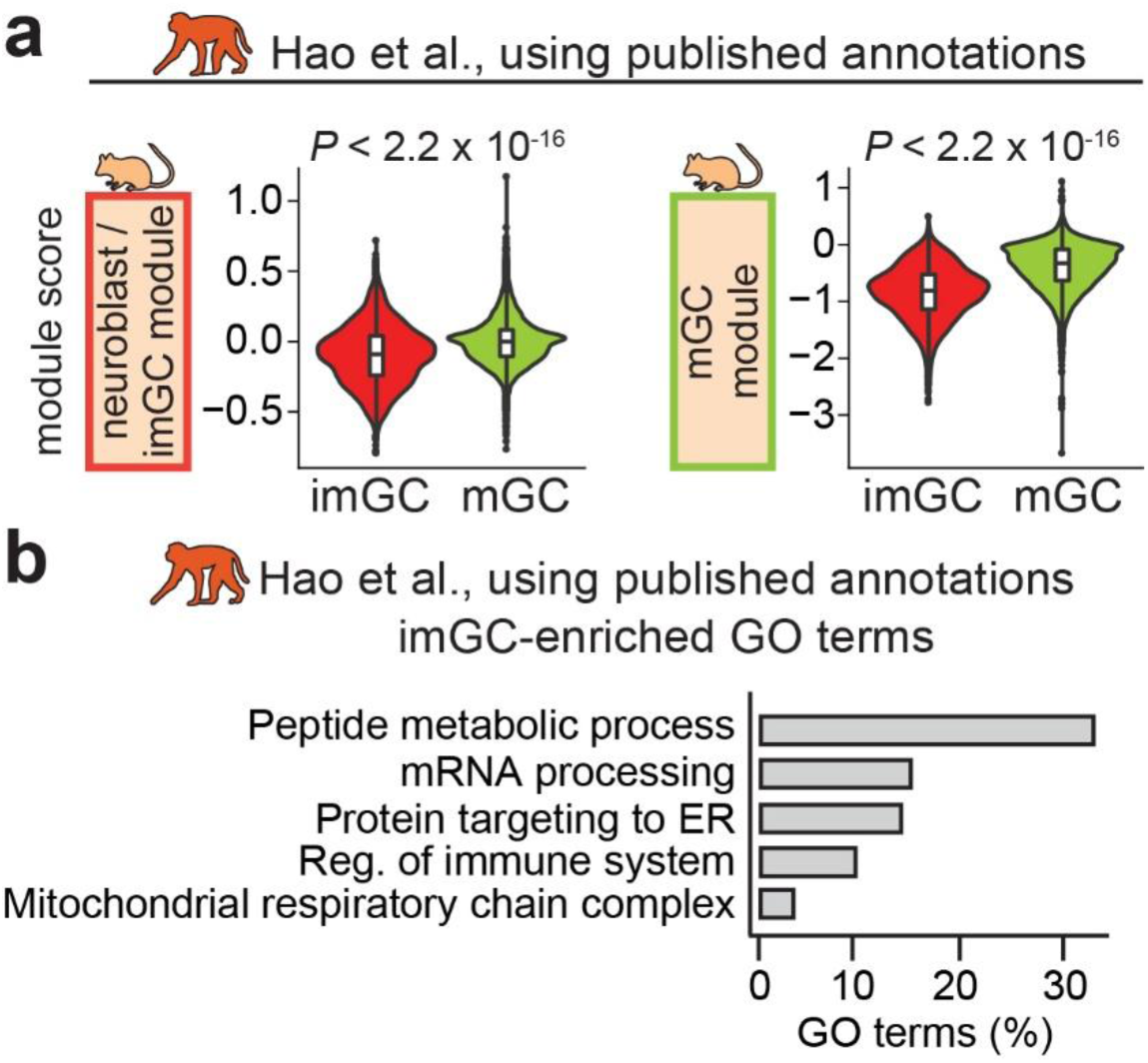
Identification and molecular characteristics of immature neurons in a published macaque hippocampal single-cell RNA sequencing dataset using traditional unsupervised clustering. **a,** Gene module analysis of the previously annotated immature dentate granule cells (imGCs) and mature dentate granule cells (mGCs) in an adult macaque hippocampus single-nucleus RNA sequencing dataset^26^ using a well-recognized published list of mouse immature progeny (neuroblast and imGC)-enriched and mGC-enriched genes^16^. **b,** Top Gene Ontology (GO) term groups for genes enriched in imGCs (using published annotations with traditional unsupervised clustering^26^). Reg., regulation.

**Extended Data Fig. 2.**
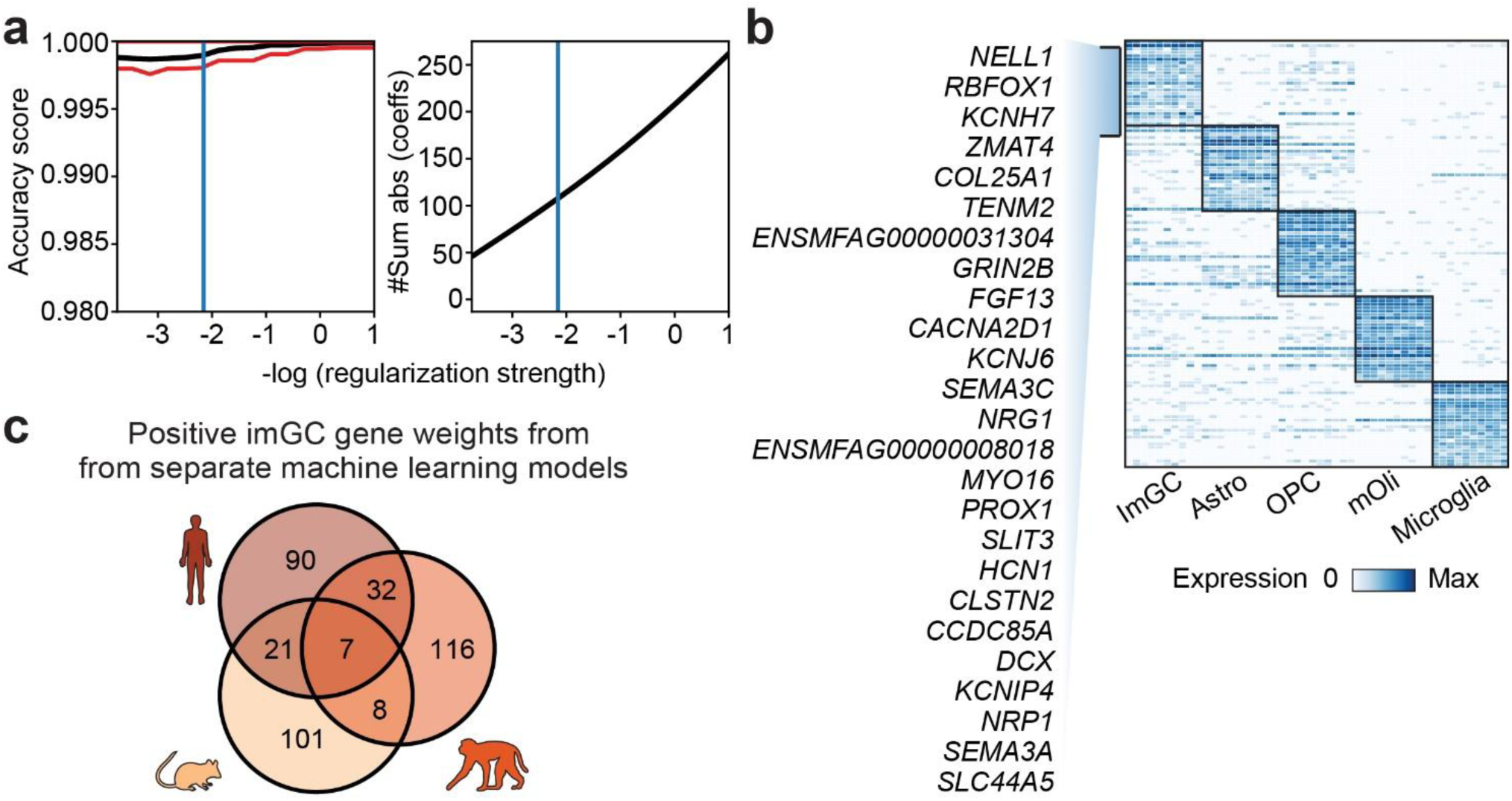
Performance of machine learning model for macaque imGCs and feature extraction and comparison of gene weights defining imGCs in different species. **a,** Measuring performance of our machine learning model for macaque datasets. Line plot showing the accuracy score of the machine learning classifier varying with decreasing regularization strength as estimated by cross-validation. Red line shows 95% confidence interval on the estimation of the accuracy score. #Sum abs (coeffs): sum of the absolute value of regression coefficients. **b,** Heatmap showing expression of top gene weights in top-scoring cells of each prototype determined by our machine learning model for macaque datasets. Genes listed are the top 25 weights defining macaque imGCs. Astro: astrocyte; OPC: oligodendrocyte progenitor cell; mOli: mature oligodendrocyte. **c,** Venn diagram of the positive gene weights defining imGCs in humans, macaques, and mice that were generated by separate machine learning models (weights for human and mouse imGCs were generated in ref^31^).

**Extended Data Fig. 3.**
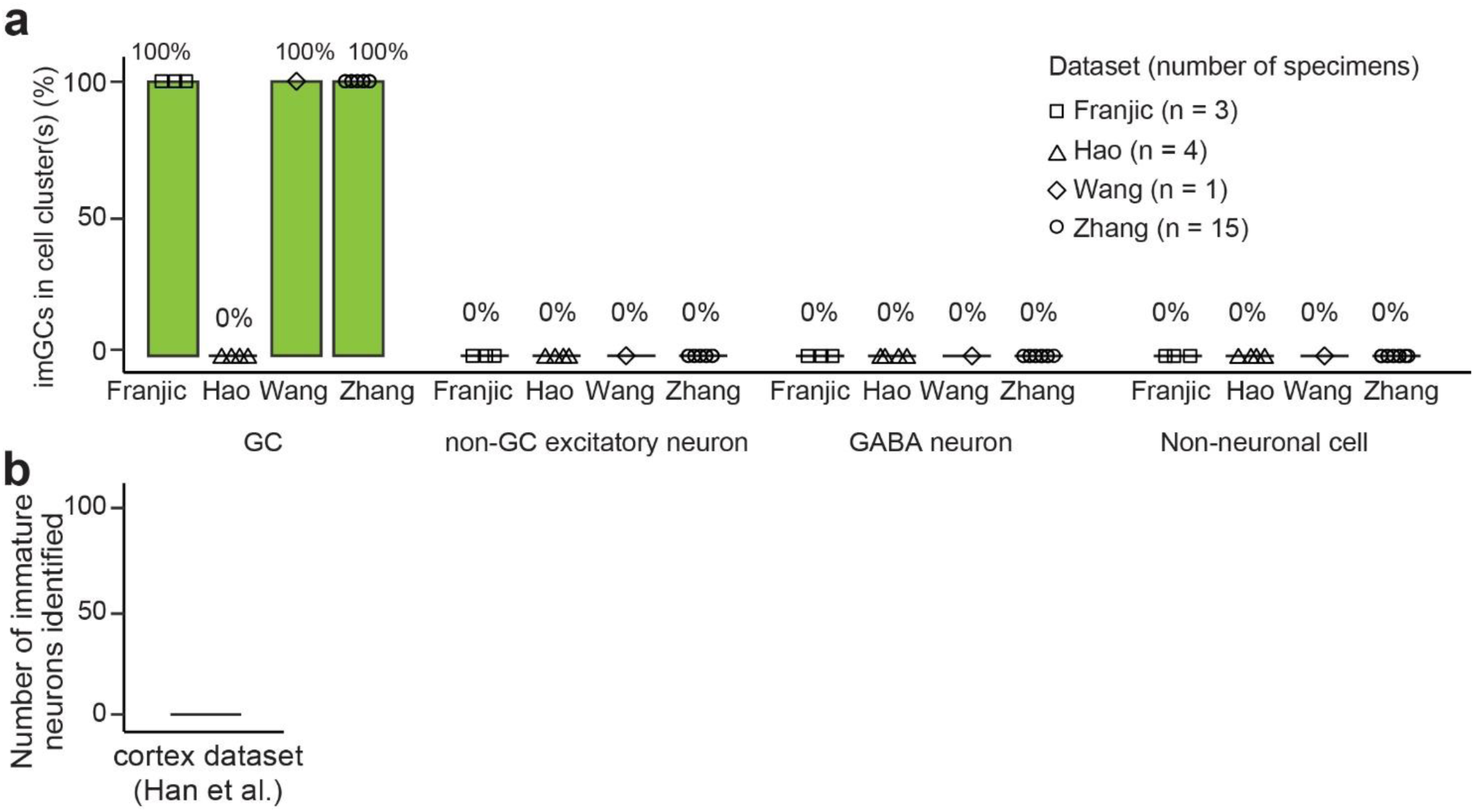
Specificity of our machine learning approach for identification of immature neurons in the macaque brain. **a,** The fractions of cells with high similarity scores (*p* ≥ 0.85) among dentate granule cell (GC), non-GC excitatory neuron, GABAergic interneuron, and non-neuronal cell clusters in various single-cell or single-nucleus RNA sequencing (scRNA-seq) datasets of the macaque hippocampus. Each dot represents data from one specimen from each study (noted by first author’s last name). Note that all imGCs identified reside in the GC clusters. **b,** No immature neurons identified using our machine learning model in an scRNA-seq dataset of two 6- year macaque neocortex (one male and one female)^54^.

**Extended Data Fig. 4.**
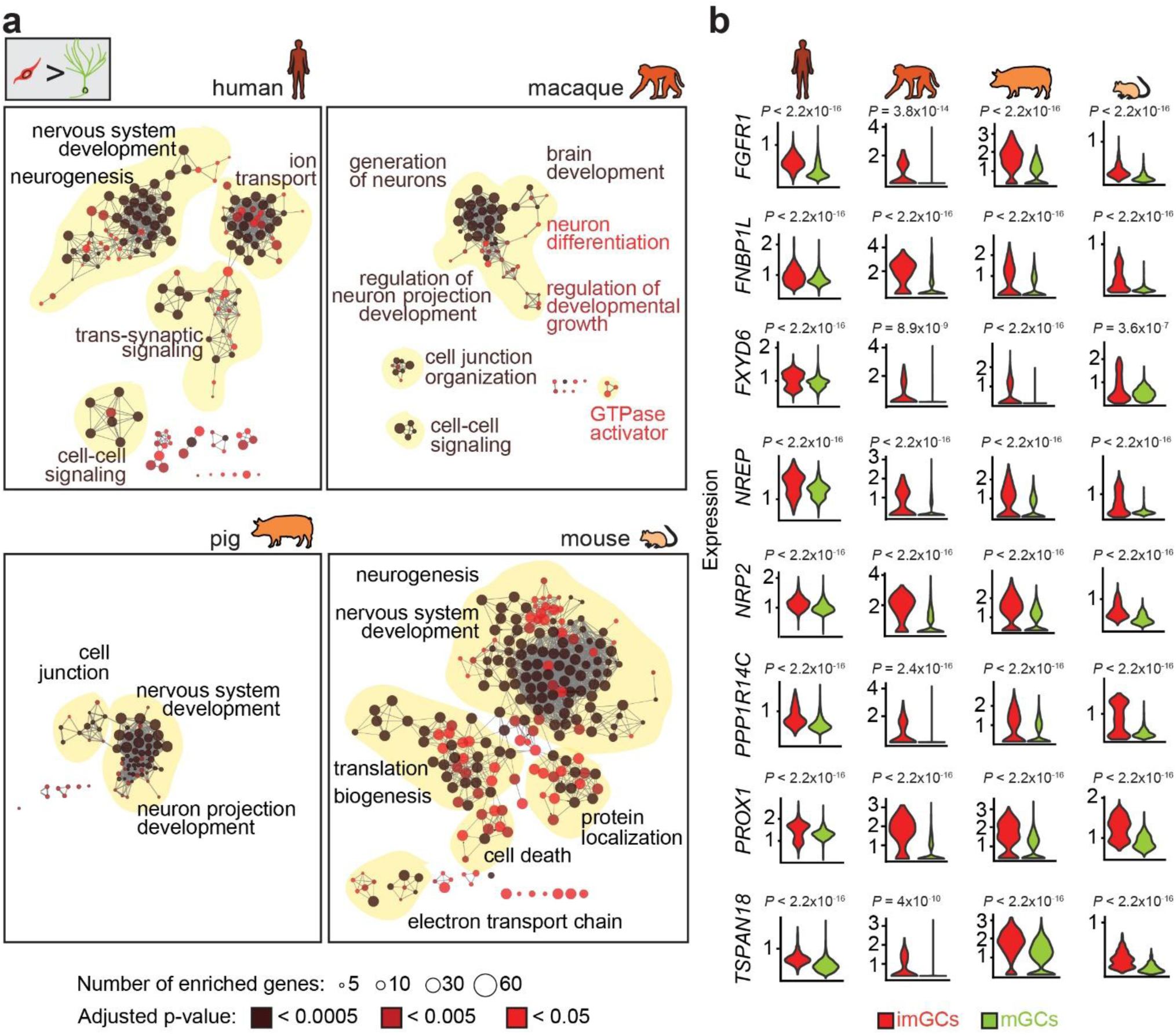
Shared molecular signatures of immature neurons in the hippocampus of different species. **a,** GO network of biological processes associated with imGC-enriched genes in different species in comparison to mGCs, coloured by FDR-adjusted p-value. Only significantly enriched nodes are displayed (one-sided hypergeometric test, FDR-adjusted *P* < 0.05). The node size represents the term enrichment significance. Examples of the most significant terms per group are shown. **b,** Violin plots showing normalized expression of 8 imGC-enriched genes shared across four species in imGCs and GCs (one-way Wilcoxon rank-sum test). A total of 9 shared genes were identified and 8 are shown here (See Fig. 3d for the plot for the 9^th^ gene, *DPYSL5*).

**Extended Data Fig. 5.**
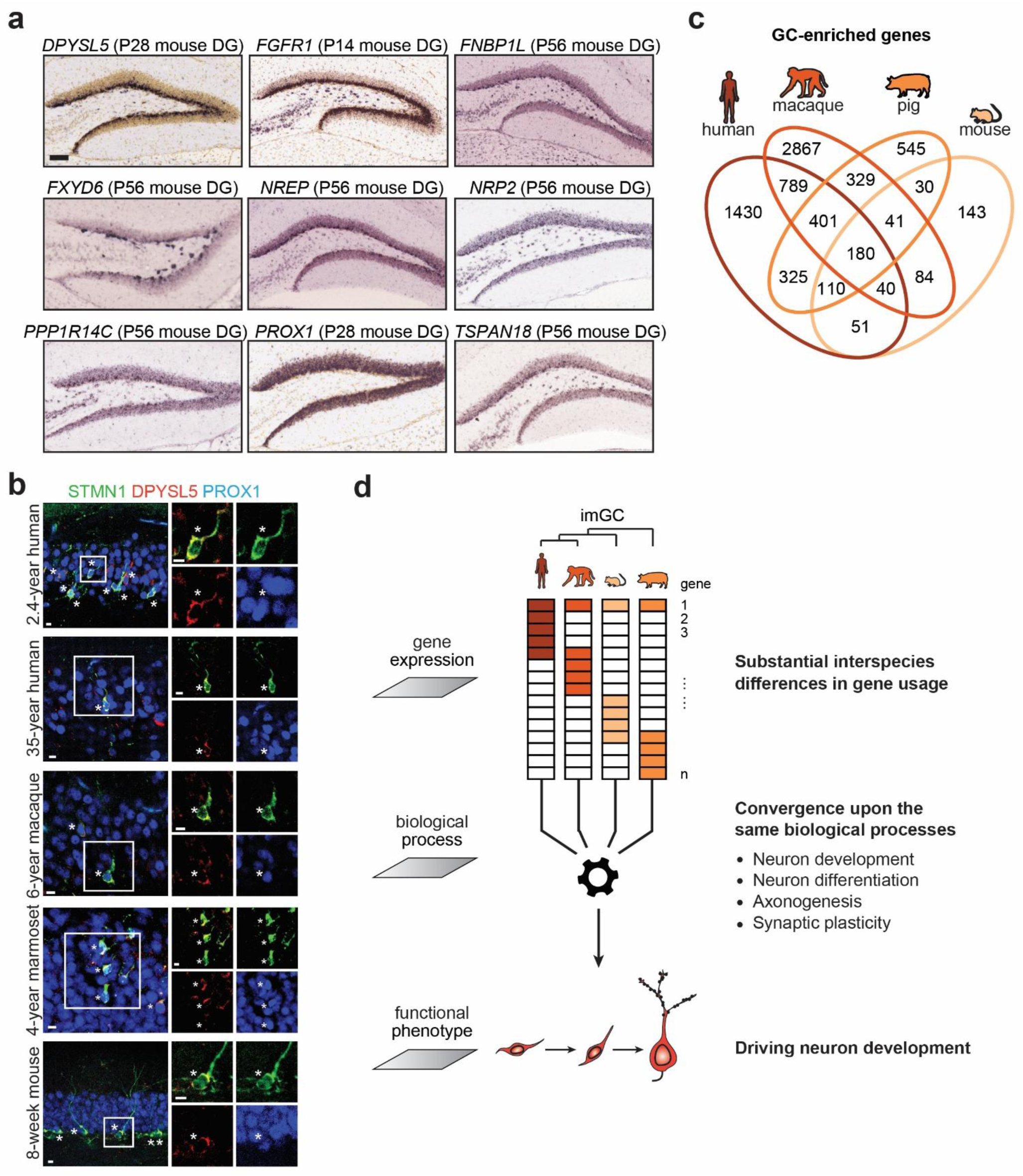
Enrichment of immature neuronal gene features in imGCs of different species. **a,** Expression patterns of 9 shared imGC-enriched genes across species in the dentate gyrus (DG) of adult mice, 8 of which show enrichment in the neurogenic subgranular zone (except for PROX1). Images are from the Allen Brain *in-situ* hybridization database^58^; https://mouse.brain-map.org/. P: postnatal day. **b,** A second set of sample confocal immunostaining images of DPYSL5 enrichment in imGCs in the hippocampi of infant and adult humans, postnatal macaques and marmosets, and adult mice. Scale bars: 10 µm. Asterisks indicate DPYSL5^+^ cells among STMN1^+^PROX1^+^ imGCs. See Fig. 3e. **c,** Venn diagram depicting the overlap of GC-enriched genes across different species when compared to other cell types within the same datasets. **d,** A schematic illustration of our working model. In contrast to the traditional concept that a single genetic variant can drive cross-species cellular innovations in immature neuron regulation, our study revealed substantial interspecies variance in highly expressed genes enriched in imGCs, which converged onto conserved biological processes, suggesting imGCs in different species may recruit and utilize species-unique molecular features to drive similar biological processes regulating neuronal development.

**Extended Data Fig. 6.**
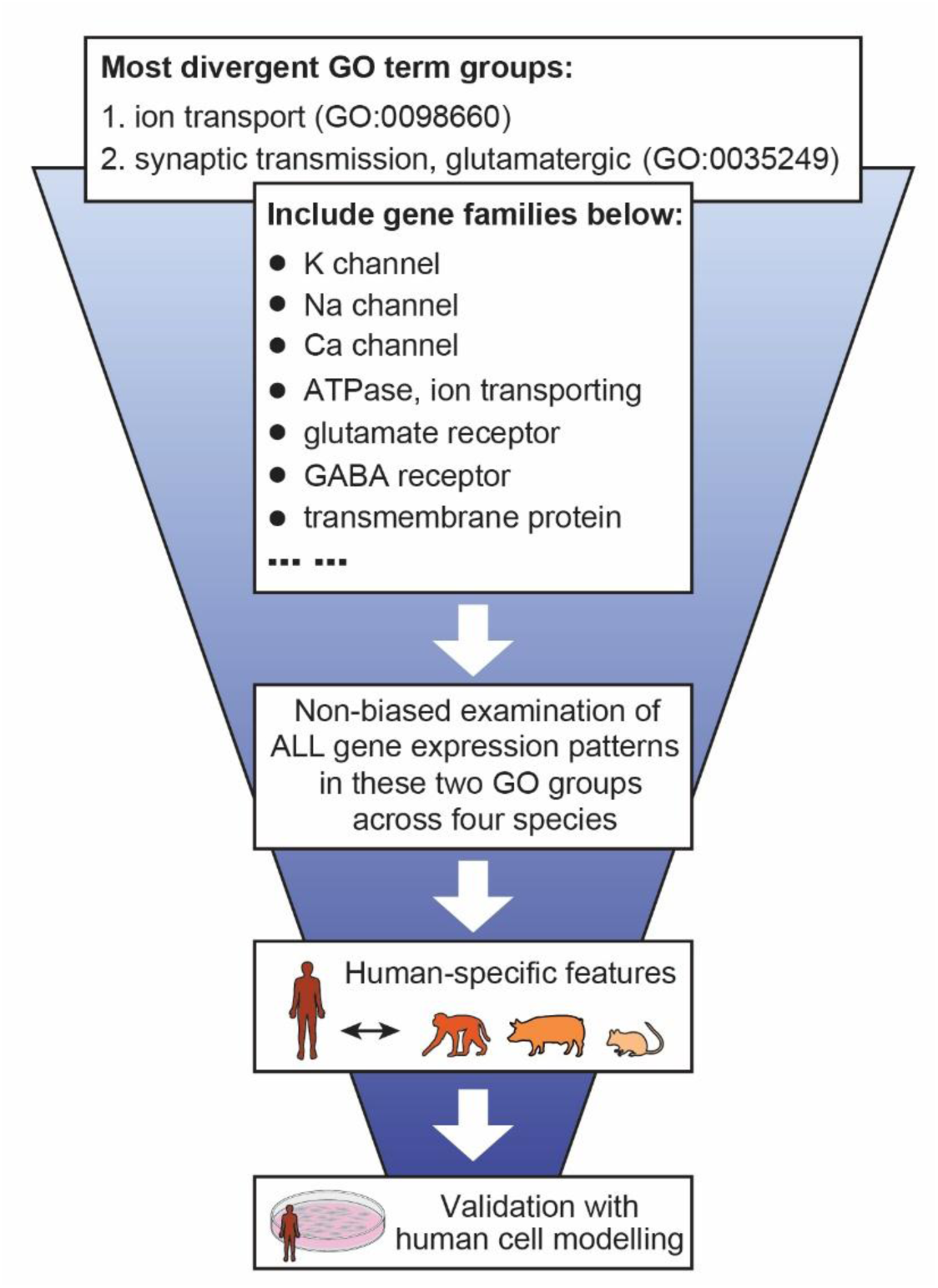
Features of imGC-enriched genes in different species. A schematic illustration of the analysis design. To explore species-specific imGC molecular features in a non-biased manner, gene expression patterns of each individual gene associated with “ion transport” and “synaptic transmission”, the two major GO terms showing the most species-specific enrichment, were plotted. See **Extended Data Fig. 7**.

**Extended Data Fig. 7.**
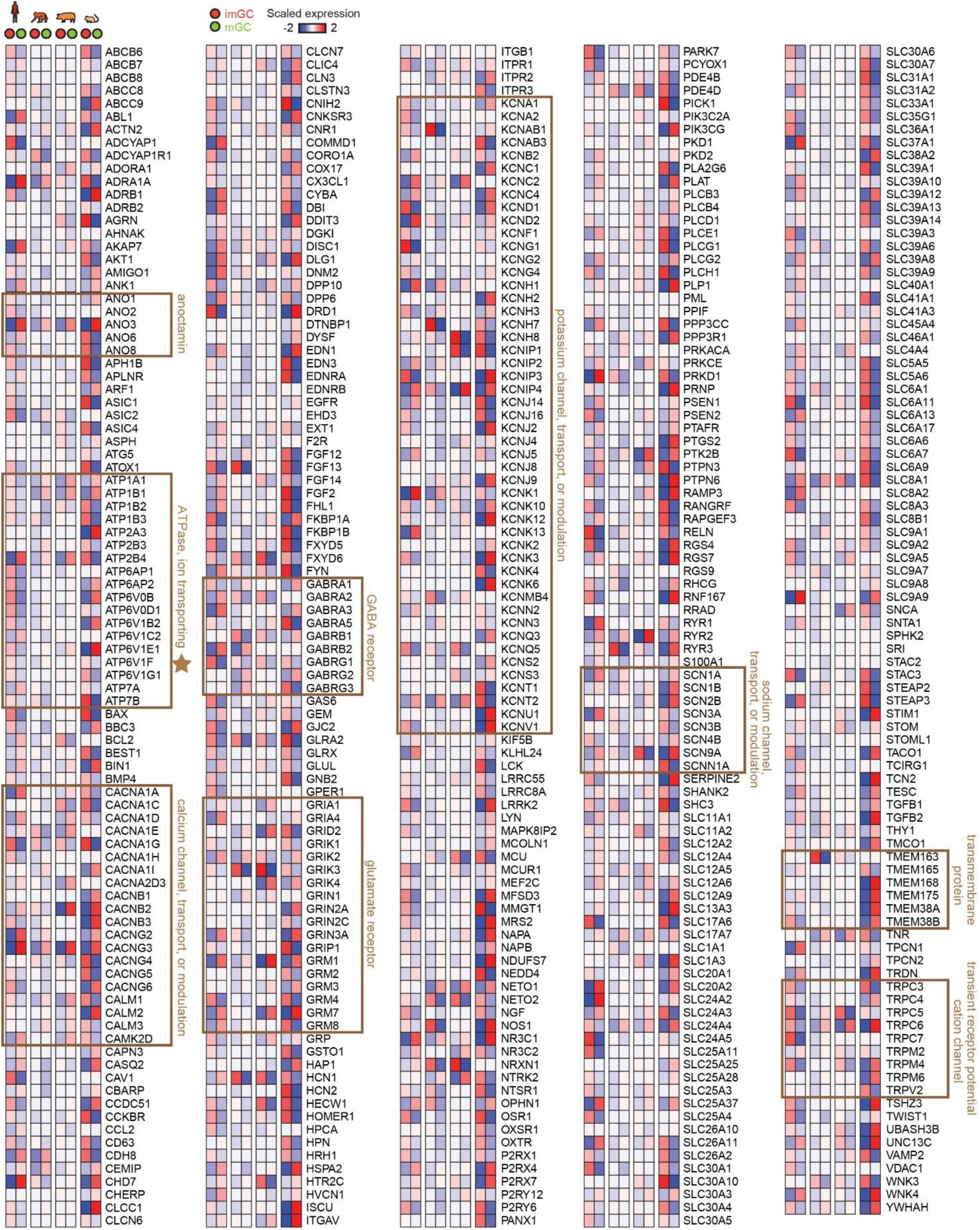
Unbiased examination of divergent imGC molecular features across four species. Red-blue heatmaps depict expression patterns of each individual gene associated with “ion transport” and “synaptic transmission”, the two major GO terms showing the most species-specific enrichment. Exemplary genes in these two categories, such as those encoding various ion channels, glutamate receptors, GABA receptors, ATPases, transmembrane proteins, among others are highlighted with square boxes. The colour bar represents z-scores of gene expression, scaled to range from −2 to 2.

**Extended Data Fig. 8.**
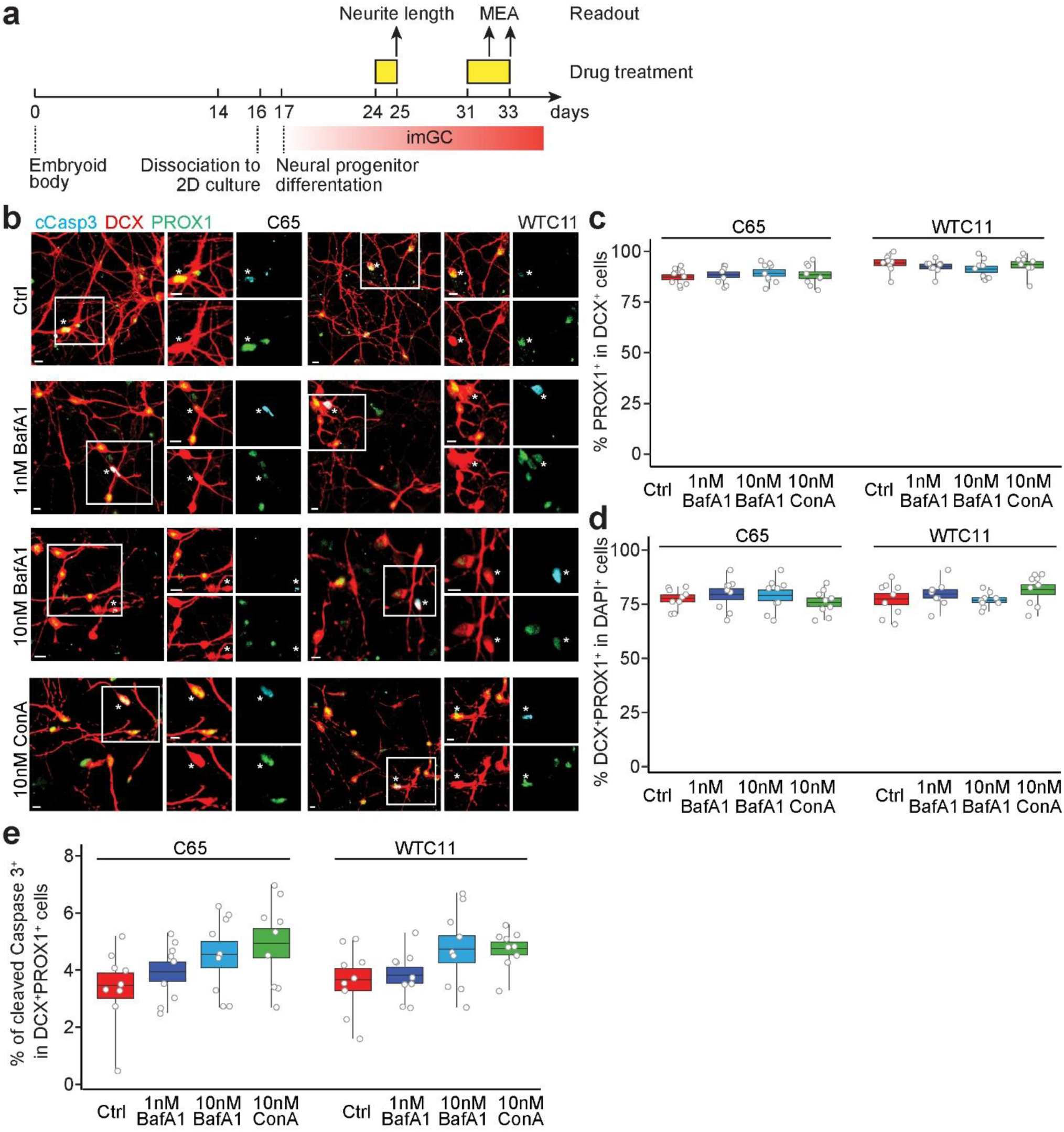
Role of lysosomal vacuolar-type H^+^-transporting ATPase in the *in vitro* human hippocampal immature neuron culture. **a,** A schematic illustration of the experimental design. Human induced pluripotent stem cell (iPSC) lines (C65 and WTC11) were differentiated into DCX^+^PROX1^+^ hippocampal imGCs prior to treatment with Bafilomycin A1 (BafA1) and Concanamycin A (ConA), two specific blockers against lysosomal vacuolar-type H^+^-transporting ATPases to measure neurite growth and neuronal activities. **b-e,** Characterization of hippocampal imGC culture derived from two independent iPSC lines. Sample confocal images (**b**) and quantification of DCX and PROX1 enrichment in the hippocampal neuron *in vitro* culture (**c, d**) and its cell death level (**e**) with different treatments. Scale bar: 10 µm. Asterisks indicate cleaved Caspase 3 (cCasp3)^+^DCX^+^PROX1^+^ imGCs (**b**). Box colours match treatment conditions. Dots represent data from individual images; the centre line represents the mean, box edges show s.e.m. and whiskers extend to the maximum and minimum values (n = 3 cultures per condition) (**c-e**). None of the quantifications were statistically significant using ANOVA post-hoc test (**c-e**).

## Supplementary Tables (in Excel files)

**Supplementary Table 1 |** Summary of published single-cell or single-nucleus RNA sequencing datasets used in the current study.

**Supplementary Table 2 |** Molecular weights defining immature dentate granule cells in macaques.

**Supplementary Table 3 |** Gene Ontology terms related to the positive gene weights defining macaque immature dentate granule cells.

**Supplementary Table 4 |** Lists of Gene Ontology terms associated with genes enriched in immature dentate granule cells across four species.

**Supplementary Table 5 |** Lists of Gene Ontology terms associated with genes uniquely enriched in immature dentate granule cells in four species.

**Supplementary Table 6 |** Summary of human specimens used for histology validation.

## Notes

### Competing Interest Statement

The authors have declared no competing interest.

